# Pediatric brain charts reveal ADHD biotypes with distinct developmental, molecular, and therapeutic profiles

**DOI:** 10.64898/2026.10.22.760978

**Authors:** Xuan Bu, Lianglong Sun, Zhao Fu, Yilu Zhao, Xiangyu Zheng, Kangfuxi Zhang, Xinyuan Liang, Zaixu Cui, Jia-Hong Gao, Gaolang Gong, Ruiwang Huang, Shuyu Li, Liang Luo, Weiwei Men, Yun Peng, Li Sun, Xiaoyi Sun, Xindi Wang, Mingrui Xia, Tengda Zhao, Litao Zhu, Xinian Zuo, Yufeng Wang, Qi Dong, Qingjiu Cao, Li Yang, Yong He

**Author notes:** Correspondence to: Yong He, PhD, Li Yang, MD, Qingjiu Cao, PhD, MD, Qi Dong, PhD. Xuan Bu and Lianglong Sun contributed to the work equally.

## Abstract

Attention-deficit/hyperactivity disorder (ADHD) exhibits profound clinical heterogeneity; however, its underlying neurobiological substrates remain unclear. Here, leveraging large-scale structural magnetic resonance imaging data from 12,663 typically developing children and 1,699 children with ADHD across Chinese and United States cohorts, we construct population-specific pediatric brain charts via hierarchical Bayesian normative modeling to quantify individual deviations in cortical morphology. Data-driven clustering identifies two robust, cross-ethnically replicable ADHD biotypes: an infra-normal biotype characterized by widespread cortical thinning and a supra-normal biotype marked by cortical thickening. Multivariate analyses reveal distinct brain–behavior architectures: cortical deviations correlate preferentially with symptom severity in the infra-normal biotype but with both symptom severity and cognitive performance in the supranormal biotype. Clinically, the infra-normal biotype demonstrates a superior response to methylphenidate over atomoxetine, whereas the supra-normal biotype responds comparably to both. Transcriptomic analyses identify biotype-specific molecular signatures converging with ADHD genome-wide association study (GWAS) risk loci but exhibiting distinct expression patterns in the brain. These findings classify ADHD heterogeneity into neurobiologically distinct, clinically informed biotypes, providing a mechanistic framework for precision psychiatry in pediatric ADHD.

## Introduction

Attention-deficit/hyperactivity disorder (ADHD) is a prevalent neurodevelopmental condition characterized by profound individual heterogeneity in clinical symptoms, cognitive profiles, and temperamental traits^1–7^. However, the neurobiological substrates of this variability remain elusive. While neuroimaging studies have identified structural^8–11^ and functional^12,13^ brain alterations in pediatric ADHD patients, conventional case control comparisons typically yield small effect sizes and inconsistent results^14^. For example, two large-scale ADHD neuroimaging investigations reported contradictory conclusions regarding cortical thickness alterations—one reporting significant group differences^10^, and the other finding no group differences^11^. Such discrepancies represent a core feature of ADHD: marked interindividual variability in brain phenotypes^15–17^.

This recognition has driven a shift toward data-driven approaches aimed at identifying neurobiologically coherent subtypes, or biotypes, to elucidate the mechanisms underlying disease heterogeneity and guide personalized interventions.

Prior efforts have delineated ADHD biotypes using structural morphology^18–20^, functional connectivity^21–24^, and multimodal features^25,26^, offering valuable insights into the neurophysiological diversity of this disorder. However, these approaches share a critical limitation: they largely overlook the dynamic neurodevelopmental context of pediatric ADHD. Children with ADHD exhibit maturational delays in cortical thickness^27^ and functional connectivity^28^, particularly within fronto-temporo-parietal networks. As a result, subtyping algorithms that fail to account for age-related changes risk grouping individuals by maturational stage rather than true neuropathological divergence. Normative modeling, or brain charting^16,29^, provides a principled solution by quantifying individual deviations from age- and sex-adjusted reference trajectories. Although a recent study applied this framework to cortical networks in ADHD^30^, its normative reference was derived from a small, retrospective, and population-biased cohort, limiting the robustness and generalizability of the resulting biotypes. Furthermore, existing ADHD biotypes largely lack mechanistic interpretation and treatment validation. To date, neuroimaging-derived biotypes have rarely been linked to differential pharmacotherapeutic responses, nor have their macroscopic brain signatures been anchored to underlying molecular mechanisms, such as spatial transcriptomic profiles. Without this multiscale integration, the proposed biotypes remain largely descriptive. Establishing replicable ADHD biotypes that are developmentally calibrated, molecularly grounded, and clinically informed therefore remains a central challenge.

Here, we address these gaps by aggregating large-scale, high-quality structural magnetic resonance imaging (MRI) data from two prospective, population-based pediatric cohorts (Chinese and U.S.), comprising 12,663 typically developing children (TDC, ages 6–19) and 1,699 children with ADHD (ages 6–17). We first constructed population-specific pediatric brain charts to quantify individual-level deviations in cortical morphology in children with ADHD. Using these individualized deviation signatures, we identified robust, reproducible ADHD biotypes and systematically evaluated their associations with clinical symptoms, cognitive function, gene expression, and treatment outcomes. Collectively, we identify two cross-ethnically replicable ADHD biotypes with distinct neurodevelopmental, cognitive, transcriptomic, and pharmacotherapeutic profiles, thereby advancing the neurobiological stratification of pediatric ADHD.

## Results

### Data samples and study design

All the data underwent rigorous quality control and harmonized analyses (Supplementary Methods). The primary discovery dataset comprised multisite structural MR data from 7,074 Chinese children across three cohorts: the Chinese Child Brain Development Project (CCBD; 6,279 TDC, 28 sites), the Peking University Sixth Hospital cohort (PKU6; 78 TDC, 505 ADHD, 1 site), and the ADHD200 cohort (130 TDC, 82 ADHD, 1 site in China) (Fig. 1A, Tables S1 and S2). The TDC samples from the PKU6 and ADHD200 cohorts were randomly split into two age-balanced halves: a training set (n = 104) for estimating site effects and a held-out testing set (n = 104) for comparisons with the ADHD group. The study workflow is outlined in Fig. 1.

**Fig. 1.**
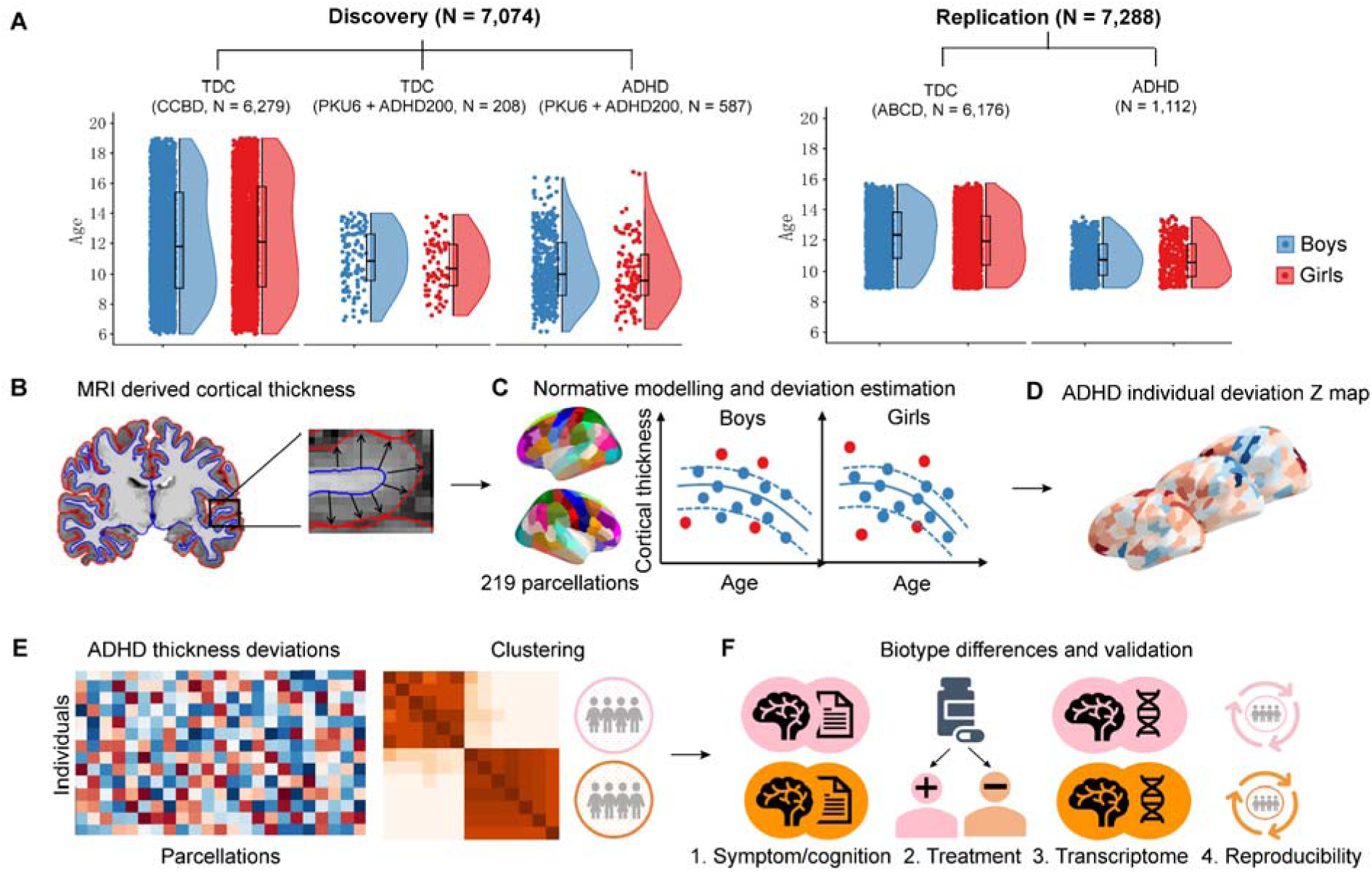
Data samples and study workflow. **(A)** Age and sex distributions across the study cohorts, displayed as box and violin plots. **(B)** Estimation of regional cortical thickness for each participant. **(C)** Construction of sex-specific normative models across 219 cortical regions. For each child with ADHD, regional thickness was compared against these normative benchmarks to quantify individual deviation profiles (Z scores). **(D)** Z maps illustrating individual cortical deviation profiles. **(E)** Clustering of children with ADHD into distinct biotypes based on their cortical deviation profiles. **(F)** Downstream evaluation of biotype-specific differences in clinical symptoms, cognitive performance, treatment response, and transcriptomic signatures, along with cross-ethnicity reproducibility.

Briefly, cortical thickness was estimated for 219 regions per participant using the Desikan–Killiany atlas^31,32^ (Fig. 1B). We constructed Chinese pediatric brain charts for these regions using a normative sample of 6,383 children (comprising the CCBD cohort and TDC training set from the PKU6 and ADHD200 datasets) (Fig. 1C). These normative models were subsequently applied to quantify regional cortical deviations from expected trajectories for each child in the testing set and each child with ADHD, yielding individualized deviation Z-maps (Fig. 1D). Spectral clustering of these deviation maps identified distinct ADHD biotypes (Fig. 1E), which were subsequently characterized across multiple domains, including symptom profiles, cognitive performance, treatment response, and spatial transcriptomic signatures (Fig. 1F). To evaluate the replicability and generalizability of our findings, we leveraged an independent validation dataset from the Adolescent Brain Cognitive Development (ABCD) study, a prospective, harmonized U.S. cohort comprising 6,176 TDC and 1,112 children with ADHD (Fig. 1A, Tables S1 and S2).

## Pediatric brain charts reveal remarkable neuroanatomical heterogeneity in ADHD

For each of the 219 cortical regions, we established Chinese pediatric normative models (n = 6,383) of cortical thickness using hierarchical Bayesian regression^33,34^, adjusting for age, sex, and scanning site. Consistent with prior literature, typically developing children exhibited widespread age-related cortical thinning across most regions (Fig. 2A). Model performance metrics are detailed in Fig. 2B and Fig. S1. Applying these normative models to children with ADHD (n = 587) and a held-out TDC testing dataset (n = 104), we generated individualized cortical deviation Z maps for each participant. Extreme deviations were defined using a threshold of |Z| > 2.0^35,36^. Notably, although no single cortical region exhibited extreme deviations in more than 15% of the ADHD cohort, more than 80% of these children had at least one region with an extreme positive or negative deviation (Fig. 2C). These results underscore the profound interindividual heterogeneity in cortical morphology among children with ADHD.

**Fig. 2.**
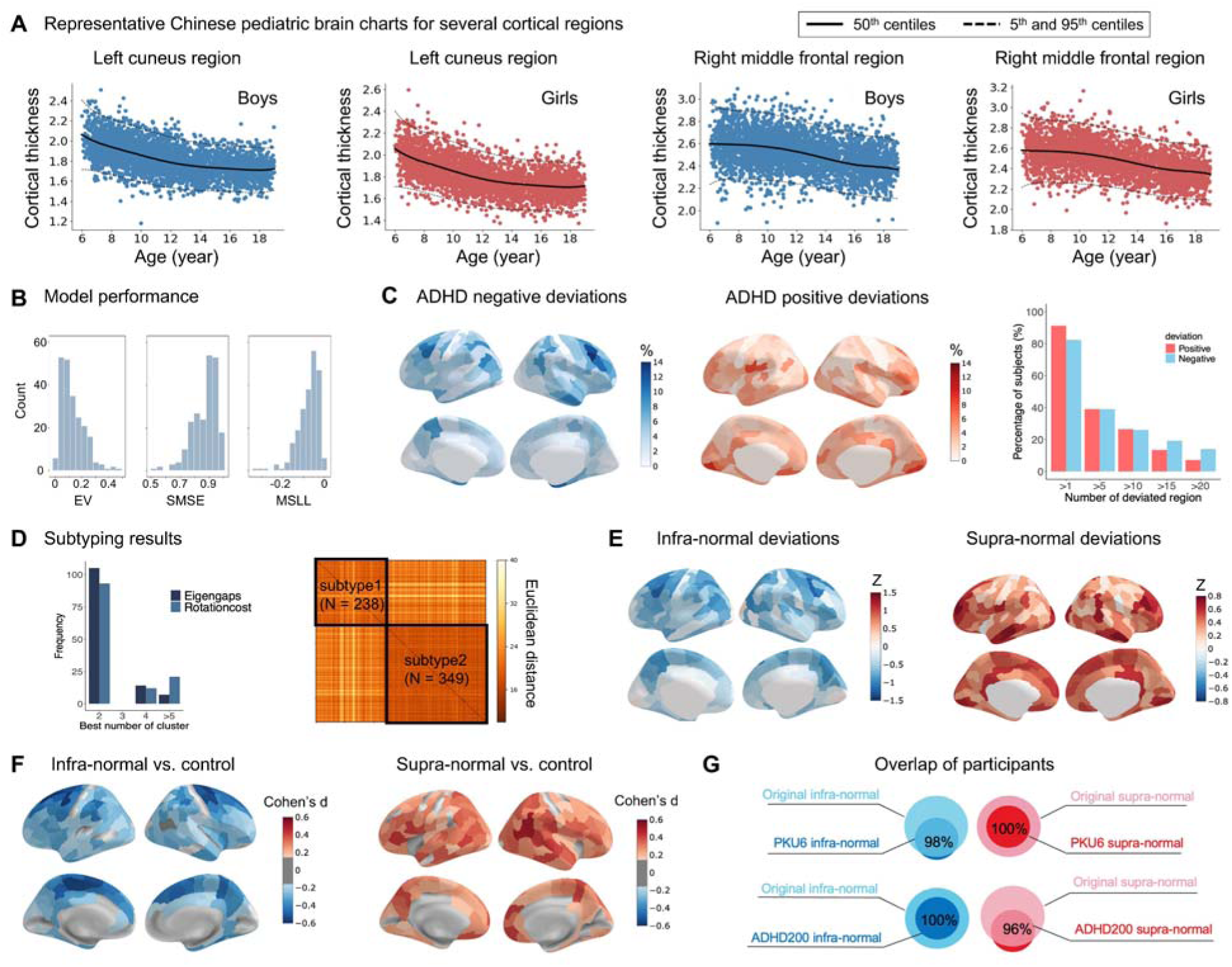
Neuroanatomical heterogeneity and identification of two ADHD biotypes using Chinese pediatric brain charts. **(A)** Representative normative trajectories (brain charts) for the left cuneus and right middle frontal cortex. **(B)** Normative model performance evaluated via explained variance (EV), standardized mean squared error (SMSE), and mean standardized log-loss (MSLL). **(C)** Spatial maps illustrating the percentage of children with ADHD exhibiting extreme cortical deviations (|Z| > 2.0) per region. The bar plot shows the percentage of children with ADHD exhibiting extreme positive (red) or negative (blue) deviations in at least one region. **(D)** Determination of the optimal number of ADHD biotypes (left) and the corresponding intersubject similarity matrix based on cortical deviation profiles (right). **(E)** Average cortical deviation maps for the infra-normal (left) and supra-normal (right) biotypes. **(F)** Case control differences in cortical deviations for the infra-normal (left) and supra-normal (right) biotypes, as assessed via one-way analysis of covariance. **(G)** Venn diagrams illustrating the overlap in biotype assignments between the independent single-site clustering and primary classification.

## Two robust ADHD biotypes characterized by individual brain deviation profiles

On the basis of individual deviation maps of cortical morphology, we computed a participant-by-participant similarity matrix among children with ADHD (n = 587). Spectral clustering applied to this matrix identified two ADHD biotypes with distinct patterns of cortical thickness deviations (Fig. 2D). Clustering stability was assessed and confirmed by bootstrap resampling across 1,000 iterations (Methods and Fig. S2). Specifically, biotype 1, termed the infra-normal subtype, exhibited widespread negative deviations, predominantly in the superior and middle frontal, precentral, and superior and medial parietal cortices (n = 238; 185 boys), whereas biotype 2, termed the supra-normal subtype, exhibited widespread positive deviations, predominantly in the prefrontal and lateral temporal cortices (n = 349; 285 boys) (Fig. 2E and Fig. S3). Case control comparisons revealed that the infra-normal biotype showed cortical thinning across extensive regions relative to TDC testing data (|Cohen’s d|: 0.15 0.64, *P*_FDR_ < 0.05), whereas the supra-normal biotype showed cortical thickening in numerous regions (|Cohen’s d|: 0.15–0.48, *P*_FDR_ < 0.05) (Fig. 2F and Fig. S4). Notably, when children with ADHD were analyzed as a single group, no significant differences in cortical thickness were detected relative to controls.

Using k-means clustering as an alternative approach, we identified two biotypes that were highly consistent with those obtained from the spectral clustering. The biotype-specific deviation patterns were strongly correlated between the two methods (Spearman’s rho = 0.99, *P* < 0.001; Fig. S5). We further repeated the biotype identification and case control analyses separately within each of the two clinical sites, PKU6 and ADHD200. Each site again yielded two ADHD biotypes with high assignment consistency relative to the original clustering: 98% overlap for the infra-normal biotype and 100% for the supra-normal biotype at PKU6 and 100% and 96%, respectively, at ADHD200 (Fig. 2G). Biotype-specific cortical thickness alterations also showed consistent spatial patterns across sites (Fig. S6). Together, these validation analyses demonstrate the robustness of the identified ADHD biotypes across clustering methods and clinical sites.

## Symptom and cognitive correlates of ADHD biotype-specific cortical deviations

We performed both univariate and multivariate analyses to examine how ADHD biotype-specific cortical deviations map onto individual differences in symptom and cognitive profiles. Although univariate analyses revealed no significant differences between the two biotypes in terms of age, sex, IQ, symptoms, or cognitive performance (Table S3), multivariate partial least-squares (PLS) regression revealed distinct patterns of covariation between brain deviations and the clinical phenotypes unique to each biotype.

For the infra-normal biotype, the first PLS component accounted for 95.8% of the variance in ADHD symptoms (*P*_perm_ = 0.004; Fig. 3A), revealing a significant correlation between the brain and symptom scores (*r* = 0.38, *P*_perm_ < 0.001; Fig. 3B). All three symptom dimensions (hyperactivity-impulsivity, inattention, and total scores) contributed to this multivariate association (Fig. 3C and Fig. S7A). Spatially, cortical deviations exhibited positive loadings in the lateral and medial parietal and lateral temporal cortices and negative loadings predominantly in the medial occipital cortex (Fig. 3D and Fig. S7B). In the supra-normal biotype, the first PLS component explained 91% of the symptom variance (*P*_perm_ = 0.02; Fig. 3E), revealing a significant correlation between the brain and symptom scores (*r* = 0.58, *P*_perm_ < 0.001; Fig. 3F). Here, positive loadings were localized mainly in the medial occipital and parietal cortices, whereas negative loadings were mapped onto the lateral frontal, temporal, and occipital cortices (Fig. 3G, H; Fig. S7C, D). Although all symptom dimensions drove the multivariate association in both biotypes, their corresponding cortical topographies differed substantially, highlighting the biotype-specific neural architectures underlying ADHD symptoms.

**Fig. 3.**
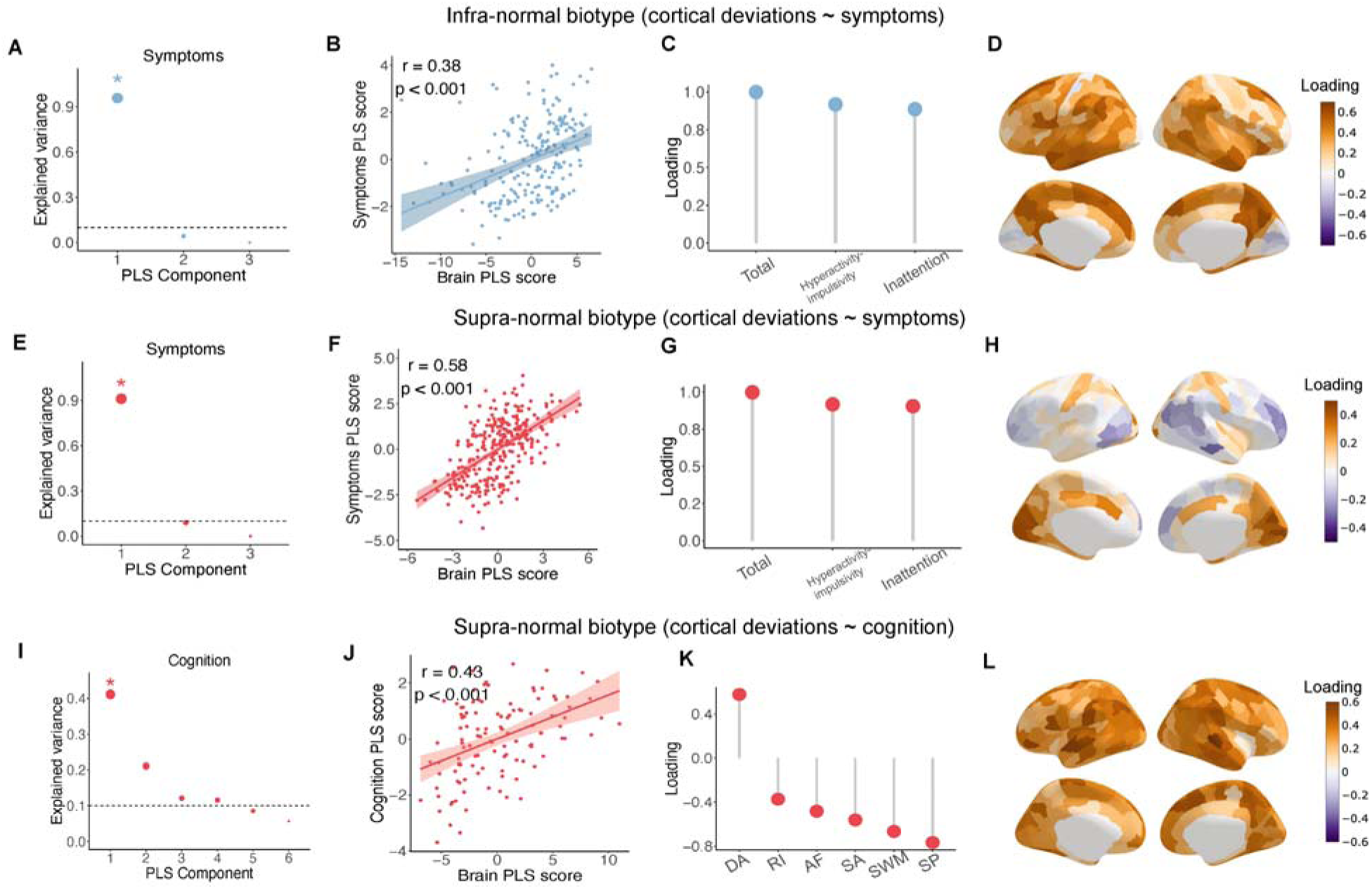
**Associations of cortical deviations with symptom and cognitive profiles across ADHD biotypes. (A–D**) Brain–symptom association derived from PLS analysis for the infra-normal biotype. **(A)** Explained variance of the PLS components. The significant component is denoted by an asterisk. **(B)** Pearson correlation between PLS brain and symptom scores. Shaded areas represent 95% confidence intervals (CIs). **(C, D)** PLS loading for individual symptom variables **(C**) and across all 219 regions (**D**). **(E–H)** Corresponding brain–symptom associations for the supra-normal biotype, formatted as in **A–D**. **(I–L)** Brain-cognition PLS association in the supra-normal biotype. Panels follow the same layout as **A–D**, displaying explained variance (**I**), brain–cognition correlations (**J**), cognitive variable loadings (**K**), and regional brain loadings (**L**). AF: attentional flexibility; SP: spatial planning; SA: sustained attention; RI: response inhibition; SWM: spatial working memory; DA: delay aversion.

With respect to cognition, the PLS analysis revealed no significant brain–cognition associations in the infra-normal biotype. In contrast, for the supra-normal biotype, the first PLS component captured 41.1% of the variance in cognition (*P*_perm_ = 0.038; Fig. 3I), indicating a significant correlation between cognition and the brain (*r* = 0.43, *P*_perm_ < 0.001; Fig. 3J). The cognitive profile was characterized by a high positive loading for delay aversion and negative loadings for spatial planning, spatial working memory, sustained attention, and attentional flexibility (Fig. 3K and Fig. S7E). The associated cortical deviations exhibited positive loadings primarily in the lateral temporal, as well as the lateral and medial parietal cortices (Fig. 3L and Fig. S7F).

## Differential treatment responses across ADHD biotypes

In the PKU6 cohort, 128 children with ADHD underwent 12 weeks of pharmacological treatment with either methylphenidate or atomoxetine. Clinical symptoms were assessed at baseline and posttreatment (Table S4). To determine whether treatment efficacy differed between medications within each biotype, we employed a two-way repeated measures analysis of variance with medication type (methylphenidate vs. atomoxetine) and time (pre- vs. posttreatment) as factors.

In the infra-normal biotype, significant medication type × time interactions were detected for the total symptom score (*F* = 5.40, *P* = 0.02; Table S5) and the hyperactivity-impulsivity score (*F* = 4.46, *P* = 0.04; Table S6) but not for the inattention score (*F* = 2.30, *P* = 0.14; Table S7). Post hoc analyses revealed that patients with the infra-normal biotype exhibited a better response to methylphenidate than to atomoxetine. This was evidenced by more pronounced reductions in both the total symptom score (methylphenidate: *F* = 51.20, *P* < 0.001; atomoxetine: *F* = 9.08, *P* = 0.009) and the hyperactivity-impulsivity score (methylphenidate: *F* = 29.52, *P* < 0.001; atomoxetine: *F* = 2.27, *P* = 0.15) (Fig. 4A, B).

**Fig. 4.**
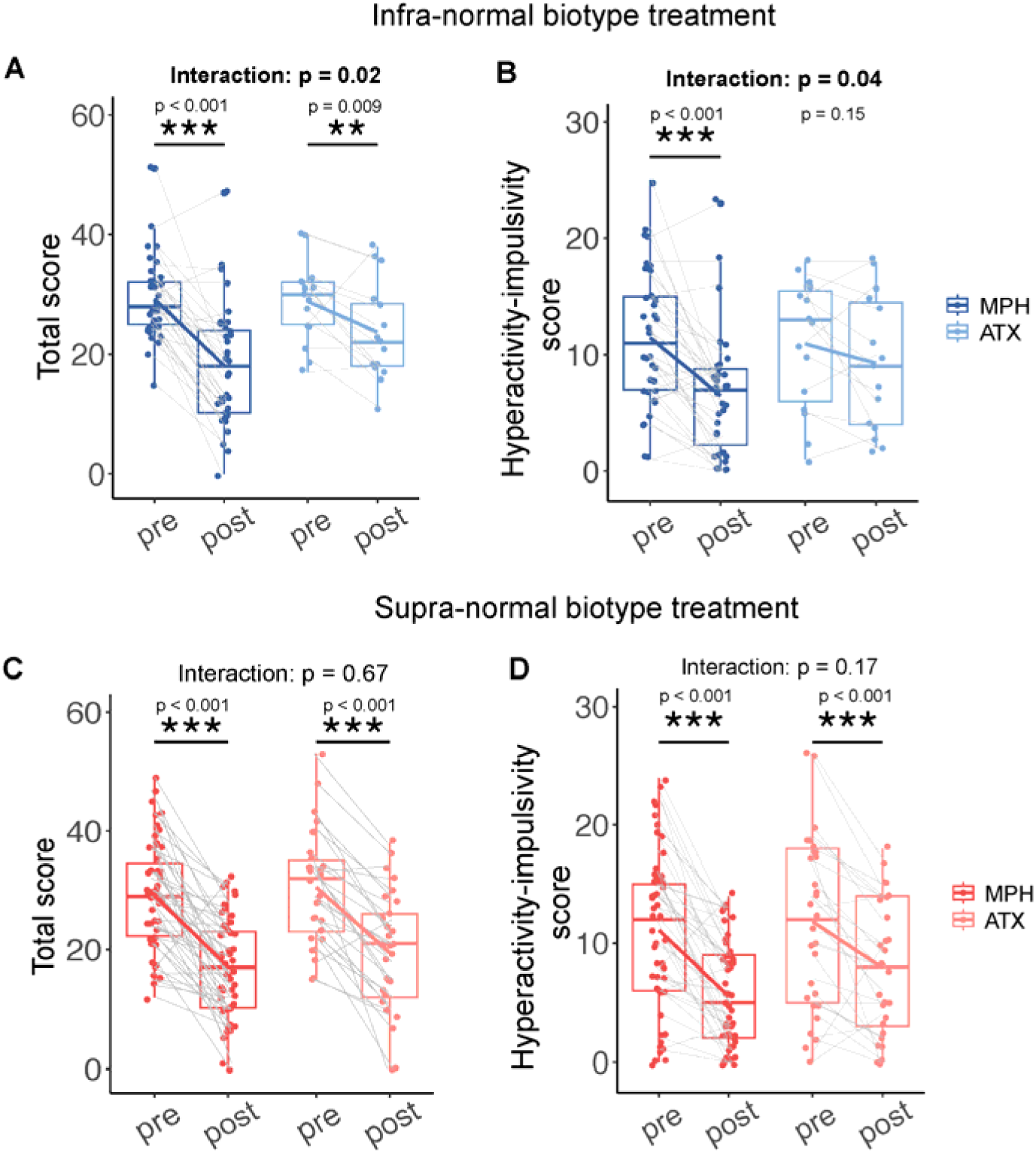
Differential treatment responses across ADHD biotypes. (A, **B)** Significant medication type × time interaction in the infra-normal biotype as assessed by total score (left panel) and hyperactivity-impulsivity score (right panel). **(C, D)** No mediation type × time interaction in the supra-normal biotype as assessed by total score (left panel) and hyperactivity-impulsivity score (right panel). MPH: methylphenidate; ATX: atomoxetine.

Conversely, for the supra-normal biotype, no significant medication type × time interactions were observed (Fig. 4C, D). Instead, a significant main effect of time was observed for the total symptom score (*F* = 127.83, *P* < 0.001; Table S5), hyperactivity-impulsivity score (*F* = 71.29, *P* < 0.001; Table S6), and inattention score (*F* = 120.25, *P* < 0.001; Table S7), indicating that both medications yielded comparable clinical benefits in this subgroup.

## Transcriptomic signatures of ADHD biotype-specific cortical deviations

To determine potential neurobiological substrates underlying cortical deviations in ADHD biotypes, we applied PLS analysis to evaluate the spatial alignment between weighted combinations of gene expression and cortical deviations in each biotype. Regional gene expression profiles were obtained from the Allen Human Brain Atlas (AHBA)^37^ and preprocessed using a standardized pipeline^38^ (Methods).

For the infra-normal biotype, the first PLS component explained 34.4% of the variance in cortical deviations (*P*_spin_ = 0.002; Fig. 5A), with a positive correlation between the PLS gene score and cortical deviations (*r* = 0.59, *P*_spin_ < 0.001; Fig. 5B). This component revealed a transcriptional profile characterized by low PLS scores mainly in the dorsal medial and lateral frontal cortices and high PLS scores in the medial and lateral occipital cortex and lateral temporal cortex (Fig. 5C). We then separately ranked the positively and negatively weighted PLS genes according to their normalized Z scores (Methods). To further understand the enrichment of PLS genes associated with specific biological processes, molecular functions, and cellular components, we performed GO enrichment analysis for the top-ranked PLS+ (1,887 genes) and PLS- genes (2,142 genes). The PLS+ genes were enriched in “chromatin remodeling”, “mRNA metabolic process”, “transcription coregulator activity”, and “transcription regulator complex” (*P*_FDR_ < 0.05), whereas the top-ranked PLS- genes were enriched in “presynapse”, “neuron projection development”, “glutamatergic synapse”, and “regulation of transsynaptic signaling” (P_FDR_ < 0.05) (Fig. 5D and Fig. S8).

**Fig. 5.**
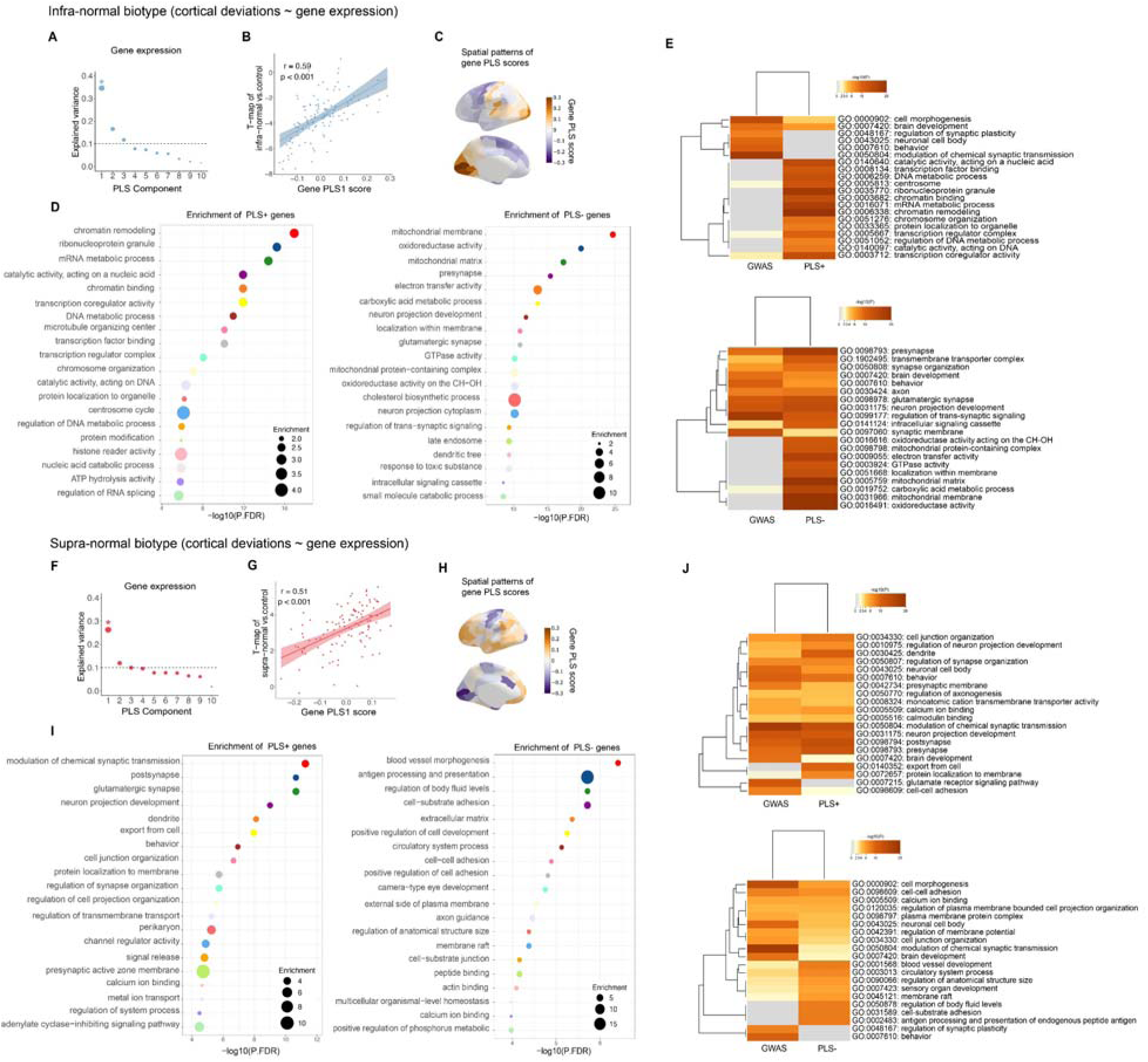
Transcriptomic associations of cortical deviations across ADHD biotypes. (A–E) Brain-gene association derived from PLS analysis for the infra-normal biotype. **(A)** Explained variance for the first 10 PLS components. The significant component is denoted by an asterisk. **(B)** Pearson correlation between regional PLS gene scores and the biotype-specific *T*-map. Shaded areas represent 95% confidence intervals (CIs). **(C)** Spatial topography of PLS gene scores across 111 left hemispheric regions. **(D)** Functional enrichment of the top-ranked PLS+ genes and PLS- genes. Node size indicates the fold enrichment of significant GO terms. **(E)** Multigene-list meta-analysis comparing infra-normal PLS genes with ADHD risk genes. The heatmap illustrates shared and specific functional enrichments, with cell color indicating statistical significance (gray: nonsignificant). **(F-J)** Corresponding brain-gene association for the supra-normal biotype, formatted as in **A–E**.

For the supra-normal biotype, the first PLS component explained 25.8% of the variance in cortical deviations (*P*_spin_ < 0.001; Fig. 5F), with a positive correlation between the PLS gene score and cortical deviations (*r* =0.51, *P*_spin_ < 0.001; Fig. 5G). High PLS scores were observed in the lateral parietal, temporal, and frontal cortices, and low PLS scores were detected in the medial visual cortex, precentral gyrus, and dorsal anterior cingulate (Fig. 5H). The top-ranked PLS+ genes (949 genes) were enriched in “modulation of chemical synaptic transmission”, “postsynapse”, “regulation of synapse organization”, and “neuron projection development” (*P*_FDR_ < 0.05), whereas the top-ranked PLS- genes (977 genes) were enriched in “blood vessel morphogenesis”, “positive regulation of cell development”, “axon guidance”, and “regulation of anatomical structure size” (*P*_FDR_ < 0.05) (Fig. 5I and Fig. S9).

To further evaluate whether these PLS-derived gene sets converge with known ADHD risk genes, we compared GO terms enriched in PLS+ and PLS- gene sets with those enriched in ADHD-associated genes identified from the latest large-scale genome-wide association study (GWAS) (Methods). A meta-analysis of multiple gene lists was performed (see the ADHD GWAS gene list in Table S8). For the infra-normal biotype, the PLS+ and GWAS gene sets shared common GO terms, including “cell morphogenesis”, “brain development”, and “transcription coregulator activity”, whereas the PLS- and GWAS gene sets shared common GO terms, including “axon”, “presynapse”, “synaptic organization”, and “neuron projection development” (Fig. 5E and Fig. S10). For the supra-normal biotype, more shared enrichment was observed. The PLS+ and GWAS gene sets shared common GO terms, including “modulation of chemical synaptic transmission”, “postsynapse”, “neuron projection development”, and “dendrite”, whereas the PLS- and GWAS gene sets shared common GO terms, including “cell morphogenesis”, “cell cell adhesion”, and “brain development” (Fig. 5J and Fig. S11). Both biotypes were enriched in genes related to neuron projection development and glutamatergic synapses but displayed distinct expression patterns in the brain.

## Biotype validation in an independent dataset

To evaluate the reproducibility and generalizability of the identified ADHD biotypes, we applied the normative modeling and subtyping framework to an independent U.S. validation dataset from the ABCD study (6,176 TDC and 1,112 children with ADHD). Using the prospective, large-scale, harmonized TDC dataset, we constructed U.S. pediatric brain charts (see model performance in Fig. S12) and quantified individual cortical deviations and biotypes. No more than 5% of the children with ADHD showed extreme deviations in any single brain region (Fig. S13), indicating a greater degree of heterogeneity in cortical morphology. Using the same clustering procedure, we also identified two ADHD biotypes (biotype 1, n = 499; biotype 2, n = 613; Fig. 6A), with biotype 1 showing predominantly negative deviations and biotype 2 showing predominantly positive deviations. The biotype-specific deviation patterns were consistent with those identified in the discovery dataset (biotype 1: *r* = 0.46, *P*_spin_ < 0.001; Fig. 6B; biotype 2: *r* = 0.32, *P*_spin_ < 0.001; Fig. 6C). Although case control differences in cortical deviations were more spatially widespread in the validation dataset (Fig. S14), their overall patterns were also consistent with those observed in the discovery dataset (biotype 1: *r* = 0.47, *P*_spin_ < 0.001; biotype 2: *r* = 0.41, *P*_spin_ < 0.001).

**Fig. 6.**
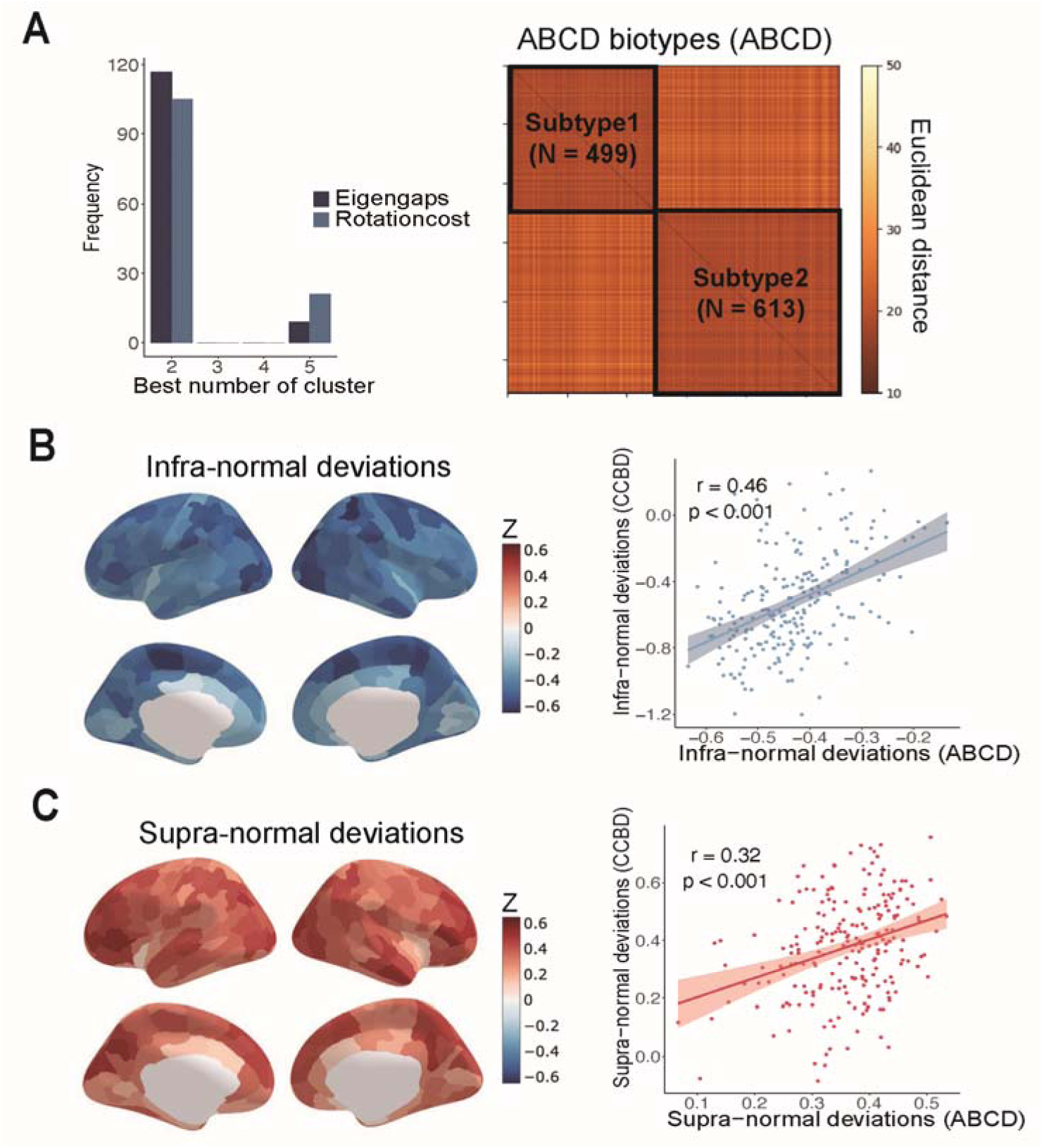
**Identification of two ADHD biotypes in the independent ABCD cohort using U.S. pediatric brain charts**. **(A)** Clustering results identifying two distinct biotypes (left panel) and the corresponding intersubject similarity matrix of cortical deviations (right panel). (**B**, **C**) Average cortical deviations for the infra-normal (**B**) and supra-normal (**C**) biotypes (left panels). Scatter plots (right panels) illustrate the spatial correlations (spin tests) of these deviation patterns between the discovery and validation cohorts.

## Discussion

Leveraging two large-scale prospective neuroimaging cohorts, we established Chinese- and U.S.-specific pediatric normative models of cortical morphology to parse the neuroanatomical heterogeneity of ADHD. This approach identified two distinct biotypes: an infra-normal biotype characterized by pronounced cortical thinning and a supra-normal biotype characterized by cortical thickening. Crucially, these two biotypes were cross-ethnically reproducible and demonstrated distinct symptomatic, cognitive, and transcriptomic profiles, as well as differential treatment responses. These findings elucidate the neurobiological basis underlying the clinical heterogeneity of ADHD and highlight the translational potential of pediatric brain chart stratification for precision psychiatry.

Previous studies of ADHD have consistently reported small effect sizes for case control differences in cortical thickness. For instance, the large-scale ENIGMA study^10^ found that cortical thinning in children with ADHD was restricted to only four regions (temporal pole, precentral, fusiform, and parahippocampal gyrus), with effect sizes ranging from -0.18 to -0.15. Similarly, the largest structural MRI study to date, utilizing the ABCD cohort^11^, detected no significant group-level differences. Consistent with these findings, our pooled case control analysis revealed no significant alterations in cortical thickness. However, moving beyond group-average comparisons, we observed widespread, individual-level deviations from normative references. The spatial distribution of these deviations varied substantially across individuals, underscoring the profound neuroanatomical heterogeneity inherent to ADHD. By stratifying patients on the basis of these individualized deviation signatures, we identified two reproducible and clinically meaningful biotypes, highlighting the critical need to parse ADHD heterogeneity through data-driven approaches.

The two ADHD biotypes exhibited opposite morphometric profiles, characterized by pronounced cortical thinning in the infra-normal biotype and cortical thickening in the supra-normal biotype. These divergent patterns align with prior reports of concurrent cortical thinning in the frontal and lateral occipitotemporal lobes^10^ and thickening in the occipital lobe^39,40^. Crucially, these biotypes mapped onto distinct symptomatic and cognitive profiles, illustrating how divergent structural architectures give rise to specific clinical phenotypes. Interestingly, the brain patterns related to symptoms were distributed in a complementary manner, with infra-normal brain being predominantly the lateral type while the supra-normal brain being mainly the medial type. The lateral temporoparietal and frontal regions are core nodes of the central executive network and the dorsal attention network which subserve working memory and top down attentional orienting. The lateral brain pattern observed in infra-biotype suggests that the symptoms may arise from weakness in the cognitive control system. On the other hand, the precuneus and medial occipital lobe are core regions of the default mode network (DMN) and the ventral visual pathway, respectively. The high contribution of these regions in supra-biotype suggests that the symptoms may be related to a failure to suppress in DMN and visual network. In line with the “modernized concept of ADHD”^41^, our results underscore the necessity of moving beyond the “average patient” paradigm. Notably, core symptoms and cognitive deficits within each biotype were associated with whole-brain deviation patterns rather than with isolated regional abnormalities. These findings implicate large-scale structural covariance networks in shaping the clinical and cognitive landscape of ADHD.

A primary clinical utility of parsing ADHD biotypes lies in advancing precision psychiatry, given that pharmacological efficacy often varies across distinct neuroanatomical profiles. This is supported by our observation that compared with atomoxetine, only the infra-normal biotype exhibited a superior therapeutic response to methylphenidate. While stimulants (methylphenidate) generally outperform nonstimulants (atomoxetine) in a subset of pediatric patients^42–44^, our results offer a neurobiological basis for stratifying this differential efficacy. The preferential responsiveness of the infra-normal biotype—characterized by widespread cortical thinning—to methylphenidate prompts mechanistic inquiry. Although both agents increase extracellular dopamine and norepinephrine levels in the prefrontal cortex via transporter blockade^45^, their broader pharmacological profiles differ. Crucially, methylphenidate additionally enhances catecholaminergic transmission within the striatum, whereas the effects of atomoxetine remain largely restricted to the prefrontal cortex^45^. The pronounced cortical thinning characteristic of the infra-normal biotype likely reflects compromised top-down cortical regulation of the basal ganglia. Consequently, methylphenidate-induced increases in dopamine and norepinephrine levels across prefrontal and mesolimbic circuits could lead to superior symptom improvements in specific infra-normal biotype.

The identified ADHD biotypes exhibited specific molecular mechanisms driving morphometric alterations. In both biotypes, morphometric deviations were spatially correlated with transcriptomic signatures enriched for neurodevelopmental processes, notably presynaptic organization, synaptic signaling, and neuron projection development. This finding aligns with evidence that synaptic disruption is a primary consequence of ADHD-associated genetic variants (CDH13, DRD4, and SLC6A3)^46–48^. Consequently, reduced neuronal and synaptic density may lead to pronounced cortical thinning in the infra-normal biotype. The pharmacological effects of stimulants also suggest that neurotransmitter dysregulation contributes to this disorder.

Furthermore, abnormalities in cell morphogenesis and axon growth (FOXP2, MEF2C, and SLC6A4)^49^ are consistent with the immaturity of cortical morphology observed in ADHD. Crucially, these transcriptomic profiles significantly overlapped with risk genes identified in the ADHD GWAS, particularly within neurodevelopmental pathways, thereby reinforcing the biological validity of our imaging-derived biotypes.

Several limitations warrant consideration. First, our cross-sectional design precludes the precise delineation of developmental trajectories; longitudinal investigations are essential to map the distinct cortical maturation profiles of these ADHD biotypes. Second, while we observed biotype-specific pharmacological effects on clinical symptoms, the extent to which these medications differentially modulate underlying brain phenotypes remains to be determined.

Third, our spatial transcriptomic analyses relied on the AHBA dataset, which maps normative gene expression in adult brains, whereas our clinical cohort included pediatric patients. Future studies leveraging postmortem transcriptomic data from pediatric ADHD cohorts will be crucial to directly link MRI-derived morphometric phenotypes with disease-specific gene dysregulation.

## Methods Participants

The present study used two independent large-scale pediatric neuroimaging datasets. The Chinese discovery dataset integrated data from the Chinese Child Brain Development (CCBD) project, the Peking University Sixth Hospital (PKU6) cohort, and the publicly available ADHD200 Study (ADHD200 cohort)^50^. The CCBD cohort is an ongoing prospective, multicenter pediatric cohort designed to characterize brain and cognitive development in Chinese school-age children. It adopts a nationwide stratified sampling framework, with a planned network of 30 data collection sites across geographically, socioeconomically and educationally diverse regions of China. The cohort comprises a cross-sectional arm covering children aged 6–18 years, which was used in the present study, and a longitudinal arm starting from 6–7 years of age. Harmonized multimodal assessments include structural and functional MRI, cognitive and behavioral measures, genetic information and environmental exposures. This design provides broad coverage of population diversity in China and supports the construction of population-specific normative references for pediatric brain development. After the application of rigorous quality control (detailed in the Supplementary methods), the final Chinese discovery sample included a total of 7,074 participants, with 6,487 typically developing children (TDC; aged 6.0–19.0 years; 3,532 boys) from the CCBD, PKU6, and ADHD200 cohorts and 587 children with ADHD (aged 6.2–16.8 years; 470 boys) from the PKU6 and ADHD200 cohorts (Fig. 1A, Table S1). Children with ADHD were diagnosed by the Schedule for Affective Disorders and Schizophrenia for School-Age Children-Present and Lifetime Version (K-SADS-PL)^51^.

The validation dataset was obtained from the Adolescent Brain and Cognitive Development (ABCD) Study^52^. The final validation sample contained 7,288 participants, including 6,176 TDC (aged 8.9–15.8 years; 3,117 boys) and 1,112 children with ADHD (aged 8.9–13.5 years; 770 boys) (Fig. 1A, Table S1).

This study was approved by the Ethics Committee of Beijing Normal University and Peking University Sixth Hospital. Written informed consent was obtained from all participants and their parents or legal guardians. Further details are provided in the Supplementary Methods.

## Image acquisition and calculation of cortical thickness

High-resolution T1-weighted structural MRI data were acquired for each dataset using 3.0 T MRI scanners (Table S2). For the discovery dataset, all participants were processed using a unified in-house CCBD structural pipeline that integrated FreeSurfer (v7.3.2). For the ABCD validation dataset, all participants were processed using the unified FreeSurfer (v6.0.1) pipeline. Briefly, for each individual, the pial surface and gray matter/white matter boundary surfaces were reconstructed. Cortical thickness was measured as the distance between linked vertices in the pial and gray matter/white matter surfaces. Using an a priori Desikan–Killiany atlas^31,32^, the mean cortical thickness was calculated for 219 regions. For details, see the Supplementary Methods.

## Constructing normative growth models of cortical morphology

To estimate site effects for the PKU6 and ADHD200 cohorts while preserving an independent TDC testing set, TDC participants from these two cohorts were randomly split into two age-balanced halves within one-year age bins. One half was combined with the CCBD TDC to form the training sample (n = 6,383; aged 6.0–19.0 years), which was used to construct regionwise normative growth models of cortical thickness for 219 cortical regions using the PCNtoolkit^53^. Briefly, cortical thickness was modeled using hierarchical Bayesian regression (HBR) as a function of age, sex and site. This framework estimates both the normative variance in cortical thickness and the uncertainty of individual predictions while flexibly modeling linear and nonlinear effects and accounting for site-related variability^33,34^. Model generalizability was assessed by 10-fold cross-validation, and regional model performance was quantified using standardized mean squared error, mean standardized log-loss and explained variance. For details, see the Supplementary Methods.

## Identifying ADHD biotypes using spectral clustering

For each individual in the TDC test and ADHD group (n = 587), the cortical thickness map was positioned onto the normative model to estimate individual deviations. For a given child with ADHD, *i*, the deviation score *Z* of the region, *j*, was calculated as follows:

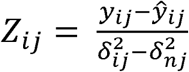

Where *y_ij_* is the true cortical thickness, *ŷ_ij_* is the predicted cortical thickness, 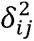 is the prediction uncertainty, and 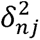 is the normative variance. Extreme positive or negative deviations were defined as |Z| >2 ^35,36^. Thus, we acquired 219 brain deviations for each child with ADHD.

To identify ADHD biotypes, we performed spectral clustering analysis on the deviation maps of 587 ADHD children using the SNFtool^54^. Briefly, the interparticipant similarity matrix (587 X 587) was estimated by calculating the Euclidean distance between the brain deviation profiles of every pair of participants. Spectral clustering was subsequently applied to the similarity matrix to cluster the participants into subgroups. The optimal number of clusters was determined according to eigengaps and rotation costs. Bootstrapping (1,000 iterations, 80% of 587 samples) was employed to validate the robustness of the subtyping. To evaluate the reproducibility of the ADHD biotypes, we 1) repeated the above analyses within each site in the discovery dataset; 2) performed normative modeling and clustering analysis in an independent validation dataset; and 3) conducted an alternative clustering method using K-means clustering. We assessed the similarity of the results across different sites, datasets, and clustering methods (details are provided in the Supplementary Methods).

We also compared cortical thickness deviations between ADHD biotypes and the TDC test using analysis of covariance (ANCOVA) with age, sex, and medication as covariates. Multiple comparisons were corrected using the false discovery rate (FDR) at *q* < 0.05.

## Relationships between cortical deviations and demographic characteristics, symptoms, and cognition among ADHD biotypes

To examine whether the ADHD biotypes exhibited differences in demographic characteristics (sex, age, and IQ), symptoms (inattention, hyperactivity-impulsivity, and total score), and cognitive function (delay aversion, sustained attention, response inhibition, spatial working memory, spatial planning, attention flexibility), we performed both univariate and multivariate analyses. The chi-square test was used to evaluate biotype differences in sex. General linear models were applied to assess biotype differences in symptoms and cognitive function, with age, sex, site, and medication used as covariates. We then used multivariate PLS regression for each biotype to explore the relationships between brain deviations (predictor variables) and symptoms/cognition (response variables). The statistical significance of the PLS components was evaluated using permutation tests (n = 5,000). For the significant PLS components, we calculated Pearson correlations between the brain score and symptom/cognition score, and the significance of the correlations was determined by permutation tests (n = 5,000). To assess the loading stability, we applied bootstrapping resampling (n = 5,000) to calculate the Z score. For details, see the Supplementary Methods.

## ADHD biotype differences in treatment response

We further investigated whether the identified ADHD biotypes responded differently to treatment. In the PKU6 cohort, 128 children with ADHD underwent 12 weeks of medication treatment with either methylphenidate or atomoxetine, and their symptoms were assessed both before and after treatment. We examined ADHD biotype differences in treatment response using a two-way (medication type, time) repeated-measures analysis of variance (ANOVA) within each biotype, with each symptom score as the dependent variable. Post hoc tests were performed to examine the effects of medication type (methylphenidate vs. atomoxetine) and time (pre- vs. posttreatment).

## Transcriptomic profiling of biotype-specific changes in cortical deviations

PLS analysis was used to evaluate the associations between biotype-specific changes in cortical deviation (*T*-map) and gene expression profiles. Human brain microarray samples were sourced from the Allen Human Brain Atlas (AHBA) (n = 6 donors)^37^. After preprocessing using the abagen toolbox^38^, the expression profiles of 8,564 genes across 111 cortical regions were obtained. The statistical significance of the PLS components was tested using spin-based spatial permutation tests^55,56^ (n = 5,000). For significant PLS components, the Pearson correlation coefficient between the gene score and the brain score was calculated, and the statistical significance was tested using spin tests (n = 5,000). We transformed the gene weight into a *Z* score by dividing by the standard error estimated from bootstrapping (n = 5,000)^57^. Using univariate one-sample *Z* tests, significant genes (FDR-corrected, *q* < 0.05) were identified and ranked separately as positive or negative weighted genes. For details, see the Supplementary Methods.

To determine the biological implications of the identified genes, we performed Gene Ontology (GO) enrichment analysis using Metascape^58^ (https://metascape.org). We selected three ontology categories, namely, biological process, molecular function, and cellular component. The significantly ranked positive and negative genes were subjected to separate analyses. To alleviate redundancy in GO terms, Metascape hierarchically clustered all the significant terms into clusters of similar terms on the basis of similarity with the kappa test score. A threshold kappa score of 0.3 was applied to split the tree into separate clusters. The most significant terms (lowest p value) within each cluster were used to represent the cluster. Finally, we investigated whether the PLS gene lists shared enrichment with risk genes for ADHD that were derived from recent GWASs^59–63^. The ADHD GWAS gene list is provided in Table S8. We performed a multigene list meta-analysis^58^ between the PLS-positive (PLS+) and PLS-negative (PLS-) gene lists and the ADHD GWAS gene list, respectively, to determine whether there were shared enrichment terms or enrichment terms selectively attributed to specific gene lists. The significance threshold for enrichment was set at an FDR *q* < 0.05.

## Author contributions

X.B., L.L.S., Q.D., Q.J.C., L.Y. and Y.H. conceived and designed the study. Q.D., Q.J.C., L.Y. and Y.H. supervised the project. X.B. and L.L.S. performed the data analysis and generated the figures. X.Y.L., X.Y.S., M.R.X. and Z.X.C. checked the accuracy of the data analysis and contributed to the interpretation of the results. Z.F., Y.L.Z., X.Y.Z, K.F.Z., L.S., Y.F.W. and L.Y. collected the data for the PKU6 cohort. J.H.G., G.L.G., R.W.H., S.Y.L., L.L., W.W.M., Y.P., X.D.W., M.R.X., T.D.Z., L.T.Z., X.N.Z, Y.H. and Q.D. contributed to the data acquisition and quality control for the CCBD cohort. X.B. wrote the first draft of the manuscript, with critical revisions from L.L.S. and Y.H. All authors reviewed the manuscript and approved the final version.

## Data availability

The ADHD200 dataset is freely available at https://fcon_1000.projects.nitrc.org/indi/adhd200/. The Allen Human Brain Atlas is available at https://human.brain-map.org/static/download. All the data necessary to replicate our results, including regional cortical thickness, cortical deviations from normative modeling, and demographic and clinical data, have been made publicly available at https://github.com/sunlianglong/ADHD-biotypes-using-cross-ethnic-pediatric-brain-charts.

## Code availability

The software packages used in this study include FreeSurfer v6.0 (https://surfer.nmr.mgh.harvard.edu/), PCNtoolkit v0.22 (https://pcntoolkit.readthedocs.io/en/latest/), SNFtool v2.3.1 (http://compbio.cs.toronto.edu/SNF/SNF/Software.html), ComBat (https://github.com/Jfortin1/ComBatHarmonization), abagen v0.1.3 (https://github.com/rmarkello/abagen), Metascape (https://metascape.org), R 4.2.2 (https://www.r-project.org), MATLAB R2021b (https://www.mathworks.com/products/matlab.html), and Python 3.8 (https://www.python.org).

The main analysis codes are available at https://github.com/sunlianglong/ADHD-biotypes-using-cross-ethnic-pediatric-brain-charts.

## Acknowledgments

This work was supported by grants from the scientific and technological innovation (STI) 2030 - the major project of the Brain Science and Brain-Inspired Intelligence Technology (2021ZD0200500 to Q.D.), the National Natural Science Foundation of China (82021004 and 82327807 to Y.H., 81671358 to L.Y., T24B2012 to L.L.S., and 824B2051 to X.Y.L.), the STI 2030-Major Projects (2022ZD0211500 to M.R.X. and 2022ZD0211300 to Z.X.C.), and the Beijing Natural Science Foundation (JQ23033 to M.R.X.). We thank the CCBD project and all its contributors for collecting the data. We thank the ABCD study for providing the data. The data used in the preparation of this article were obtained from the Adolescent Brain Cognitive Development Study (https://abcdstudy.org), which is held in the NIMH Data Archive. This is a multisite, longitudinal study designed to recruit more than 10,000 children aged 9–10 and follow them for more than 10 years into early adulthood. The ABCD Study® is supported by the National Institutes of Health and additional federal partners under award numbers U01DA041048, U01DA050989, U01DA051016, U01DA041022, U01DA051018, U01DA051037, U01DA050987, U01DA041174, U01DA041106, U01DA041117, U01DA041028, U01DA041134, U01DA050988, U01DA051039, U01DA041156, U01DA041025, U01DA041120, U01DA051038, U01DA041148, U01DA041093, U01DA041089, U24DA041123, and U24DA041147. A full list of supporters is available at https://abcdstudy.org/federal-partners.html. A list of the participating sites and a complete list of the study investigators can be found at https://abcdstudy.org/consortium_members/. ABCD consortium investigators designed and implemented the study and/or provided data but did not necessarily participate in the analysis or writing of this report. This manuscript reflects the views of the authors and may not reflect the opinions or views of the NIH or ABCD consortium investigators. The ABCD data repository grows and changes over time. Finally, we thank all participating children and their parents or legal guardians.

