## Supplementary Info for "Pediatric brain charts reveal ADHD biotypes with distinct developmental, molecular, and therapeutic profiles"

### Supplementary methods

#### 1. Participants

##### 1.1 Chinese cohorts

###### CCBD cohort

The Chinese Child Brain Development (CCBD) study is an ongoing, prospective, multicenter pediatric cohort designed to characterize brain development in Chinese children and adolescents. This project aims to acquire harmonized multimodal MRI (structural MRI, diffusion MRI, resting-state fMRI, and task fMRI), alongside comprehensive behavioral, cognitive, environmental, and genomic data from more than 30,000 participants aged 6–18 years. The CCBD incorporates both cross-sectional (ages 6–18) and longitudinal (ages 6–7 at baseline) components. By utilizing a nationwide stratified sampling framework, the cross-sectional cohort was recruited through a network of 30 data collection sites distributed across geographically, socioeconomically, and educationally diverse regions. For the present study, only structural MRI data from the cross-sectional cohort (n = 6,456) were utilized. The exclusion criteria included cognitive impairments, a history of neuropsychiatric disorders, traumatic brain injury, severe physical illness, psychoactive medication use, substance abuse, and MRI contraindications.

###### PKU6 cohort

This dataset included 566 children with ADHD (age range: 6.2–16.8 years; 450 boys) and 88 typically developing children (TDC) (age range: 6.8–13.7 years; 54 boys). Participants with ADHD were recruited from the Peking University Sixth Hospital. Clinical diagnoses were made by senior child psychiatrists according to the Diagnostic and Statistical Manual of Mental Disorders, Fourth Edition (DSM-IV) criteria and were further confirmed using the Schedule for Affective Disorders and Schizophrenia for School-Age Children-Present and Lifetime Version (K-SADS-PL). Symptom severity was assessed using the ADHD Rating Scale-IV [1]. The exclusion criteria for the ADHD cohort included (1) a history of neurological illness, significant head trauma, or severe physical illness; (2) left-handedness; (3) a history of other major psychiatric disorders or intellectual disability; and (4) a full-scale intelligence quotient (IQ) less than 80, as measured by the Wechsler Intelligence Scale for Chinese Children-Revised [2]. TDC participants were recruited from local primary and middle schools, adhering to identical exclusion criteria with the additional requirement of no lifetime history of ADHD.

A subset of 128 children with ADHD from the PKU6 cohort underwent a 12-week longitudinal follow-up to evaluate treatment response. Participants received pharmacotherapy with either methylphenidate (n = 84) or atomoxetine (n = 44). The initial dose was 18 mg/day for methylphenidate and 10 mg/day for atomoxetine. Dosages were titrated weekly on the basis of clinical efficacy and adverse effects. The maximum daily dose was 54 mg/day for methylphenidate and 100 mg/day or 1.2 mg/kg for atomoxetine.

###### ADHD200 cohort

This dataset included 86 children with ADHD (age range: 8.3–14.0 years; 75 boys) and 133 TDC (age range: 8.1–14.0 years; 78 boys) obtained from the Peking University (PKU) site of the ADHD200 consortium. To ensure age and ethnic comparability with the other two Chinese cohorts, only participants under 14 years of age from the PKU site were included. Clinical diagnoses were initially evaluated using the Computerized Diagnostic Interview Schedule IV and subsequently confirmed via the K-SADS-PL. Symptom severity was assessed using the ADHD Rating Scale-IV, and IQ was measured with the Wechsler Intelligence Scale for Chinese Children-Revised. The exclusion criteria included left-handedness, an IQ less than 80, head trauma, neurological illness, a history of major psychiatric disorders, and substance abuse. The ADHD group included both medication-free and medicated children with ADHD; for the latter, psychostimulant medications were withdrawn 24–48 hours prior to MRI acquisition.

##### 1.2 United States cohort

**Adolescent Brain Cognitive Development (ABCD)**

The ABCD study is an ongoing multisite, longitudinal cohort tracking more than 10,000 children across the United States [3]. Participants were aged 9–11 years at baseline and were primarily recruited via probability sampling of public and private elementary schools across 21 sites. This project aims to acquire harmonized multimodal MRI (structural MRI, diffusion MRI, resting-state fMRI, and task-state fMRI). The exclusion criteria included major medical or neurological conditions, a history of traumatic brain injury, and contraindications to MRI. Demographic and neuroimaging data were obtained from ABCD Data Release 5.1. Only structural MRI data were used in this study. ADHD diagnoses were made using the validated, parent-reported computerized Kiddie Schedule of Affective Disorders and Schizophrenia for DSM-5 (KSADS-COMP). The inclusion criteria for the ADHD group included current ADHD (ksads_14_853_p = 1), ADHD-NOS (ksads_14_856_p =1) and ADHD in partial remission (ksads_14_855_p = 1). Individuals with comorbid intellectual disability, bipolar disorder, or schizophrenia spectrum disorders were further excluded, in accordance with previously established criteria [4]. TDC were defined as the absence of any lifetime psychiatric disorders, as confirmed by both parent- and children's self-reported K-SADS-COMP.

Given that the baseline data cover only a restricted age range of 9–11 years, we incorporated longitudinal MRI scans spanning ages 9–16 years to construct US normative models over a broader developmental period. To ensure broader age representation while retaining only a single scan per participant, we implemented an age-stratified sampling strategy. Specifically, the 9–16-year range was divided into 1-year age bins, and scans were iteratively selected from each participant's longitudinal data, prioritizing bins with the smallest sample sizes. This procedure yielded a final cross-sectional, age-balanced sample comprising children with ADHD aged 9–14 years (drawn from baseline and 2-year follow-up) and TDC aged 9–16 years (drawn from baseline, 2-year, and 4-year follow-ups). In total, 8,086 TDC and 1,361 children with ADHD were included in the analyses (Table S1).

#### 2. Clinical and cognitive assessment

##### 2.1 ADHD symptoms

ADHD symptoms were quantitatively evaluated by the ADHD Rating Scale-IV. This scale contains 18 items corresponding to the 18 symptoms in the DSM-IV. The scale is used to quantify ADHD symptoms on a 4-point frequency scale; the answer options are 0 = never, 1 = sometimes, 2 = often, and 3 = always. The scale includes a hyperactivity-impulsivity score, an inattention score, and a total score.

##### 2.2 Assessment of cognition

Cognitive functions in participants with ADHD were assessed using the Cambridge Neuropsychological Test Automated Battery (CANTAB), a well-validated computerized test battery (https://cambridgecognition.com/cognitive-function). Detailed descriptions of the specific tests, corresponding outcome measures, and the cognitive domains evaluated in this study are summarized below.

| Test | Measurement | Cognition Domain |
|---|---|---|
| Cambridge Gambling Test | Delay aversion | Delay aversion |
| Rapid Visual Information Processing | Probability of a hit | Sustained attention |
| Stop Signal Task | Stop-signal reaction time | Response inhibition |
| Spatial Working Memory | Between errors | Spatial working memory |
| Stockings of Cambridge | Problems solved | Spatial planning |
| Intradimensional/Extradimensional Shifts | Stages completed | Attention flexibility and maintenance |

#### 3. MRI acquisition

All the structural MRI data were obtained using 3.0-T MRI scanners. The detailed scanning parameters of each dataset and site are listed in Table S2.

#### 4. Imaging data quality control (QC)

##### 4.1 Chinese cohorts (CCBD, PKU6, and ADHD200 cohorts)

Structural MRI data from the CCBD cohort (n = 6,456) underwent rigorous quality control, including visual inspection to ensure (1) no obvious structural abnormalities or imaging artifacts (e.g., Gibbs ringing); (2) successful brain extraction; (3) accurate anterior commissure–posterior commissure (AC-PC) alignment; and (4) successful brain tissue segmentation and cortical surface reconstruction. Following rigorous QC, a total of 6,279 TDC were retained for subsequent analyses (Table S1).

For the PKU6 and ADHD200 cohorts, raw structural MRI data were visually checked using a multipoint review process in accordance with the Human Connectome Project Structural QC guidelines 5. Each image was assigned a rating score on a 5-point scale (0 = bad, 1 = poor, 2 = fair, 3 = good, 4 = excellent). Only images with a score greater than 2 were advanced to FreeSurfer processing. Postreconstruction QC was performed according to the ENIGMA protocol (http://enigma.ini.usc.edu/protocols/imaging-protocols/) to ensure that cortical surface reconstructions corresponded to known anatomical boundaries. On the basis of these criteria, 505 children with ADHD and 78 TDC were retained from the PKU cohort (excluding 61 children with ADHD and 10 TDC), and 82 children with ADHD and 130 TDC were retained from the ADHD200 cohort (excluding 4 children with ADHD and 3 TDC). In total, 587 children with ADHD and 208 TDC from these two clinical cohorts were included in the final analyses (Table S1).

##### 4.2 US cohort (ABCD study)

Structural MRI data were excluded on the basis of the QC criteria recommended by the ABCD consortium (https://wiki.abcdstudy.org/release-notes/imaging/quality-control.html), which included (1) abnormal structural images, as indicated by “mrif_score” values of 0 (“Image artifacts prevent radiology read”) or 4 (“Consider immediate clinical referral”); (2) failure to pass initial QC measures, as indicated by an “iqc_t1_ok_ser” value of 0; (3) failure to pass Freesurfer-based quality assurance variables, as indicated by a QC score (“fsqc_qc”) of 0 (“reject”) or responses of >1 (“moderate” to “severe”) on motion (“fsqc_qu_motion”), pial overestimation (“fsqc_qu_pialover”), white matter underestimation (“fsqc_qu_wmunder”), and inhomogeneity (“fsqc_qu_inhomogeneity”); and (4) lack of recommendation for T1w inclusion, as indicated by a score of 0 on “imgincl_t1w_include”. These procedures resulted in the exclusion of data from 141 children with ADHD and 413 TDC. The final sample included 6,176 TDC and 1,112 children with ADHD; detailed demographic characteristics are summarized in Table S1.

#### 5. Processing of structural MRI

For the discovery dataset, structural MRI data were processed using a unified, in-house CCBD structural pipeline integrating FreeSurfer (v7.3.2). Specifically, the processing steps included intensity normalization, skull stripping, spatial normalization to the Talairach space, cortical segmentation, tessellation of the gray matter (GM)/white matter (WM) boundary, and automated topological defect correction. The reconstructed surfaces were constrained to a spherical topology and corrected for geometrical and topological abnormalities. Vertexwise cortical thickness was calculated as the shortest distance between the GM/WM boundary and the pial surface. The Desikan–Killiany anatomical atlas was utilized to subdivide 68 cortical regions into 219 distinct parcellations [6], from which the mean cortical thickness for each region was extracted. For the ABCD validation dataset, structural data were processed using the unified FreeSurfer (v6.0.1) pipeline with an identical procedure.

#### 6. Normative modeling and model performance

For the discovery dataset, we constructed a normative model of cortical thickness for each of the 219 brain regions using a training sample of 6,383 TDC (age range: 6.0–19.0 years). These models estimate the normative range of regional cortical thickness on the basis of a set of covariates (age, sex, and scanning site). To mitigate nuisance variance introduced by multisite data—which cannot be attributed to interindividual biological heterogeneity—we employed hierarchical Bayesian regression (HBR) [7]. This approach models sex and site effects by incorporating random slopes, intercepts, and noise. Within a partial-pooling framework, HBR imposes shared prior distributions over sex- and site-specific parameters. This assumes that while parameters may vary across sexes and sites, they are drawn from a common underlying distribution. Consequently, these shared priors regularize the model parameters, preventing overfitting to small batches from any specific sex or site. Normative modeling was implemented using the PCNtoolkit (https://pcntoolkit.readthedocs.io/en/latest/) [8]. Region-wise, sex-specific trajectories are available at https://github.com/sunlianglong/ADHD-biotypes-using-cross-ethnic-pediatric-brain-charts.

To assess model generalizability, we employed a 10-fold cross-validation strategy. The TDC training data were split into ten folds; the model was iteratively trained on nine folds, whereas the remaining fold was withheld to evaluate the generalization performance. This process was repeated 10 times, ensuring that each fold served as the test set exactly once. Model performance for each brain region was quantified using the explained variance (EV), the standardized mean squared error (SMSE), and the mean standardized log-loss (MSLL). These normative models were subsequently applied to data from Chinese children with ADHD (n = 587) and a hold-out TDC test dataset (n = 104),

For the validation dataset (the ABCD cohort), an identical analytic strategy was applied. The normative models were trained on data from 3,262 TDC and were subsequently tested on 1,112 children with ADHD and a hold-out test set of 2,914 TDC, with age, sex, site, and race as covariates.

**7. Spectral clustering and clustering stability**

The similarity matrix of cortical deviations across children with ADHD was calculated using a Euclidean distance with a nearest neighbor (K) value of 30 and a normalization parameter (α) of 0.8, which resulted in a 587×587 similarity matrix. The similarity (W) between patients $x_i$ and $x_j$ is computed. We denote $\rho\ (x_i, x_j)$ as the Euclidean distance between patients $x_i$ and $x_j$. We then use a scaled exponential similarity kernel to determine the weight of the edge as follows:

$$W(i,j) = \exp(-\frac{\rho^2\ (x_i, x_j)}{\mu\varepsilon_{i,j}})$$

where μ is a hyperparameter that can be empirically set and $\varepsilon_{i,j}$ is used to eliminate the scaling problem. Here, we define $\varepsilon_{i,j}$ as follows:

$$\varepsilon_{i,j} = \frac{mean(\rho(x_i, N_i) + mean(\rho(x_j, N_j)) + \rho(x_i, x_j)}{3}$$

where $mean(\rho(,))$ is the average value of the distances between $x_i$ and each of its neighbors. More detailed methodology is outlined in [9]. To find the optimal values of the parameters and number of clusters, we performed a grid search using the ranges K=10–30 and α=0.3–0.8, as recommended by the SNFtool developers. For each iteration, the number of clusters was determined using the function *estimateNumberOfClustersGivenGraph*. Features with higher normalized mutual information (NMI) values had greater contributions to clustering. As a result, the two clusters were consistently optimal on the basis of both eigengaps and the rotation cost. Spectral clustering (function *spectralClustering*) was subsequently applied to the similarity matrix across 1000 iterations to resample 80% of the subjects. A silhouette plot was generated to illustrate the similarity of the subjects within each subgroup to the subjects in the other subgroups.

To evaluate clustering stability, we conducted 1,000 bootstrap resamplings of 80% of the subjects. For each iteration, we calculated the following: (1) the percentage of time that each subject clustered with every other subject and (2) the adjusted Rand index (ranging from 0 to 1), which measures the overlap between two clusters (a value of 0 means that completely random clusters overlap, and a value of 1 means that the clusters are identical).

To evaluate the cluster reproducibility, we also performed the above analyses within each site (PKU6 and ADHD200). Once clusters were obtained from each site, we first examined whether subjects overlapped between single-site biotypes and the original biotypes. Afterward, for each site, we compared cortical thickness differences among ADHD biotypes and controls using analysis of covariance (ANCOVA) with age, sex, and medication as covariates. Multiple comparisons were corrected using the false discovery rate (FDR) at $q < 0.05$.

##### 8. Symptom and cognitive correlates of ADHD biotype-specific cortical deviations

We employed partial least squares (PLS) analysis to examine the relationship between cortical deviation and symptoms/cognition in each ADHD biotype. PLS is a multivariate latent variable model that captures associations across two types of data (i.e., brain and behavior). The PLS model finds pairs of brain and behavioral weights such that the linear combination of the brain and behavioral variables maximizes the covariance between the resulting pairs of latent components. Given two matrices, $X_{n\times p}$ and $Y_{n\times q}$, where n is the number of participants and p and q are the numbers of behavioral (X) and brain (Y) variables, respectively, PLS finds u and v, which are loading vectors that maximize cov (Xu, Yv). In our analysis, the symptom variables included total symptom score, inattention score, and hyperactivity-impulsivity score, which were available for 508 participants (infra-normal biotype: 211; supra-normal biotype: 297). The cognitive variables included delay aversion, probability of hit, stop-signal reaction time, number of errors, number of problems solved, and number of stages completed; these variables were available for 164 participants (infra-normal biotype: 49, supra-normal biotype: 115).

A permutation test (5,000 times) was applied to test the significance of the PLS component, where the brain matrix was fixed and the rows of the behavioral matrix were shuffled to eliminate linkages between brain variables and symptoms/cognitive variables. To evaluate the stability of the contribution loading of each variable, we performed bootstrap resampling 5,000 times. Since random sampling can induce axis reflection, which causes a sign change in the loadings, or axis rotation, which changes the ordering of components, we matched the order and sign of the resampled loadings to the original ones. The Z score for each brain loading and symptom/cognitive loading was then calculated by dividing the standard error estimated from bootstrapping.

##### 9. Transcriptomic signatures of ADHD biotype-specific cortical deviations

###### 9.1 Regional gene expression analysis

The Allen Human Brain Atlas (AHBA) (http://www.brain-map.org) [10] provides human brain microarray samples from 3,702 brain regions. The samples were collected from six neurotypical adult brains (age = 42.50 ± 13.38 years; male/female = 5/1). Since only two brains had right hemisphere samples, we used only microarray data from the left hemisphere in our study. The AHBA data were preprocessed according to updated standardized workflows using the abagen toolbox (Supplementary Methods) [11]. Briefly, the steps included 1) updating probe-to-gene annotations; 2) filtering probes on the basis of expression intensity (probes with an intensity less than the background in ≥50.00% of samples across donors were discarded); 3) selecting probes with the most consistent pattern of regional variation across donors on the basis of differential stability; 4) assigning samples to the brain atlas used; 5) normalizing tissue sample expression across genes using a robust sigmoid function; and 6) averaging samples assigned to the same brain region for each donor and then across donors. Finally, a matrix of 111 regions × 8,564 genes was constructed.

The detailed preprocessing of the microarray data was performed as follows. The AHBA data were processed with the abagen toolbox (version 0.1.3; https://github.com/rmarkello/abagen) using a 111-region volumetric atlas in Montreal Neurological Institute (MNI) space. First, the microarray probes were reannotated using data provided by Arnatkeviciute et al. [12]. The probes not matching a valid Entrez ID were discarded. Next, the probes were filtered according to their expression intensity relative to background noise such that the probes with intensities less than the background in ≥50.00% of samples across donors were discarded. When multiple probes indexed the expression of the same gene, we used the probe with the most consistent pattern of regional variation (highest differential stability) across donors. The differential stability was calculated as follows:

$$DS\ (p) = \frac{1}{\binom{N}{2}} \sum_{i=1}^{N-1} \sum_{j=i+1}^{N} \rho[B_i(p), B_j(\mathrm{p})]$$

where ρ is the Spearman's rank correlation of the expression of a single probe (p) across regions in two donors $B_i$ and $B_j$ and N is the total number of donors. Here, regions correspond to the structural designations provided in the ontology from the AHBA. The MNI coordinates of the tissue samples were subsequently updated to match those generated via nonlinear registration via advanced normalization tools (ANTs; https://github.com/chrisfilo/alleninf). Samples were assigned to brain regions in the provided atlas if their MNI coordinates were within 2 mm of a given parcel. To reduce the potential for misassignment, sample-to-region matching was constrained by hemisphere and gross structural divisions (i.e., cortex, subcortex/brainstem, and cerebellum) such that, e.g., a sample in the left cortex could be assigned only to an atlas parcel in the left cortex. All tissue samples not assigned to a brain region in the provided atlas were discarded. Afterward, a robust sigmoid function [13] was used to normalize the tissue sample expression values across genes as follows:

$$x_{norm} = \frac{1}{1 + \exp(-\frac{x - \langle x \rangle}{IQR_x})}$$

where $\langle x \rangle$ is the median and $IQR_x$ is the normalized interquartile range of the expression of a single tissue sample across genes. Afterward, the normalized expression values were rescaled to the unit interval as follows:

$$x_{scaled} = \frac{x_{norm} - \min(x_{norm})}{\max(x_{norm}) - \min(x_{norm})}$$

Gene expression values were then normalized across tissue samples using an identical procedure. Samples assigned to the same brain region were averaged separately for each donor and then across donors. Finally, genes whose differential stability was greater than 0.1 were retained since genes with high differential stability are related to biological relevance, disease, drug targets and literature citations [14]. The processing yielded a regional expression matrix of 111× 8,564.

##### 9.2 Null model for the spin test

Spatial autocorrelation-preserving permutation tests (spin tests) [15] were used to test the significance of associations between gene expression and cortical deviations (https://github.com/frantisekvasa/rotate_parcellation). The spin test generates randomly rotated spherical projections that preserve the spatial contiguity of the empirical brain map. First, we obtained spherical coordinates of regional centroids from the FreeSurfer annotation file for the brain parcellation used (111 brain regions from left hemisphere). Afterward, the regional coordinates were randomly rotated 5,000 times to generate a null distribution. The p value was calculated as the proportion of null explained variance that was greater than the empirical explained variance.

#### Supplementary results

#### Table S1. Demographics of participants in the datasets

| Dataset | Site | Group/Number | Age (years) | Sex (F/M) | Ethnicity/Race | #ADHD presentations (ADHD-C, ADHD-H/I, and ADHD-I) |
|---|---|---|---|---|---|---|
| Discovery | PKU6 | TDC (n = 78) | 10.06 (1.95) | 50/28 | Asian | NA |
| | | ADHD (n = 505) | 10.09 (2.13) | 399/106 | Asian | 64/6/94 |
| | ADHD200-PKU | TDC (n = 130) | 11.23 (1.69) | 78/52 | Asian | NA |
| | | ADHD (n = 82) | 11.61 (1.70) | 71/11 | Asian | 34/1/47 |
| | CCBD-site01 | TDC (n = 50) | 14.58 (3.31) | 31/19 | Asian | NA |
| | CCBD-site02 | TDC (n = 85) | 11.98 (3.25) | 42/43 | Asian | NA |
| | CCBD-site03 | TDC (n = 310) | 12.56 (3.60) | 168/142 | Asian | NA |
| | CCBD-site04 | TDC (n =184) | 12.63 (3.59) | 101/83 | Asian | NA |
| | CCBD-site05 | TDC (n = 70) | 11.71 (3.77) | 33/37 | Asian | NA |
| | CCBD-site06 | TDC (n = 289) | 12.98 (3.96) | 146/143 | Asian | NA |
| | CCBD-site7 | TDC (n = 157) | 12.51 (3.53) | 74/83 | Asian | NA |
| | CCBD-site8 | TDC (n = 88) | 14.98 (3.51) | 51/37 | Asian | NA |
| | CCBD-site9 | TDC (n = 60) | 11.31 (3.45) | 35/25 | Asian | NA |
| | CCBD-site10 | TDC (n = 87) | 11.79 (3.78) | 46/41 | Asian | NA |
| | CCBD-site11 | TDC (n = 72) | 11.55 (3.44) | 35/37 | Asian | NA |
| | CCBD-site12 | TDC (n = 105) | 12.39 (3.63) | 52/53 | Asian | NA |
| | CCBD-site13 | TDC (n = 97) | 10.11 (2.96) | 39/58 | Asian | NA |
| | CCBD-site14 | TDC (n = 113) | 10.21 (2.94) | 55/58 | Asian | NA |

| | | | | | | |
|---|---|---|---|---|---|---|
| | CCBD-site15 | TDC (n = 115) | 13.34 (3.43) | 67/48 | Asian | NA |
| | CCBD-site16 | TDC (n = 106) | 11.36 (3.41) | 54/52 | Asian | NA |
| | CCBD-site17 | TDC (n = 122) | 11.94 (3.51) | 61/61 | Asian | NA |
| | CCBD-site18 | TDC (n = 68) | 10.96 (3.08) | 41/27 | Asian | NA |
| | CCBD-site19 | TDC (n = 184) | 12.53 (3.76) | 100/84 | Asian | NA |
| | CCBD-site20 | TDC (n = 94) | 11.57 (3.85) | 49/45 | Asian | NA |
| | CCBD-site21 | TDC (n = 115) | 11.88 (3.85) | 59/56 | Asian | NA |
| | CCBD-site22 | TDC (n = 428) | 12.33 (3.78) | 223/205 | Asian | NA |
| | CCBD-site23 | TDC (n = 151) | 12.31 (3.85) | 80/71 | Asian | NA |
| | CCBD-site24 | TDC (n = 58) | 12.62 (4.04) | 34/24 | Asian | NA |
| | CCBD-site25 | TDC (n = 78) | 12.39 (3.92) | 39/39 | Asian | NA |
| | CCBD-site26 | TDC (n = 187) | 12.38 (3.77) | 102/85 | Asian | NA |
| | CCBD-site27 | TDC (n = 130) | 11.85 (3.58) | 64/66 | Asian | NA |
| | CCBD-site28 | TDC (n = 291) | 11.03 (3.24) | 146/145 | Asian | NA |
| Validation | ABCD-site01 | TDC (n = 99) | 11.56 (1.68) | 49/50 | 26 Hispanic/24 White/3 Asian/2 Black/44 Other | NA |
| | | ADHD (n = 1) | 13.00 (-) | 1/0 | 1 Hispanic/0 White/0 Asian/0 Black/0 Other | NA |
| | ABCD-site02 | TDC (n = 299) | 12.55 (1.90) | 181/118 | 20 Hispanic/197 White/2 Asian/4 Black/76 Other | NA |
| | | ADHD (n = 16) | 11.92 (1.76) | 8/8 | 1 Hispanic/7 White/0 Asian/0 Black/8 Other | NA |
| | ABCD-site03 | TDC (n = 356) | 11.66 (2.01) | 178/178 | 222 Hispanic/37 White/2 Asian/24 Black/71 Other | NA |
| | | ADHD (n = 43) | 11.09 (1.78) | 22/21 | 23 Hispanic/3 White/0 Asian/5 Black/12 Other | NA |
| | ABCD-site04 | TDC (n = 315) | 12.18 (1.92) | 171/144 | 35 Hispanic/169 White/4 Asian/26 Black/81 Other | NA |
| | | ADHD (n = 80) | 11.61 (1.82) | 42/38 | 10 Hispanic/36 White/1 Asian/6 Black/27 Other | NA |

| | | | | | |
|---|---|---|---|---|---|
| ABCD-site05 | TDC (n = 170) | 11.89 (1.94) | 89/81 | 12 Hispanic/99 White/3 Asian/30 Black/26 Other | NA |
| | ADHD (n = 41) | 11.45 (1.80) | 26/15 | 2 Hispanic/21 White/1 Asian/9 Black/8 Other | NA |
| ABCD-site06 | TDC (n = 286) | 12.19 (1.98) | 153/133 | 30 Hispanic/156 White/8 Asian/17 Black/75 Other | NA |
| | ADHD (n = 51) | 11.63 (1.89) | 27/24 | 4 Hispanic/23 White/0 Asian/4 Black/20 Other | NA |
| ABCD-site07 | TDC (n = 88) | 11.24 (1.61) | 48/40 | 8 Hispanic/43 White/1 Asian/12 Black/24 Other | NA |
| | ADHD (n = 43) | 11.00 (1.64) | 28/15 | 4 Hispanic/18 White/0 Asian/10 Black/11 Other | NA |
| ABCD-site08 | TDC (n = 124) | 11.87 (2.02) | 63/61 | 17 Hispanic/47 White/23 Asian/1 Black/36 Other | NA |
| | ADHD (n = 21) | 11.31 (2.04) | 12/9 | 2 Hispanic/5 White/1 Asian/1 Black/12 Other | NA |
| ABCD-site09 | TDC (n = 165) | 11.48 (1.99) | 86/79 | 37 Hispanic/54 White/4 Asian/8 Black/62 Other | NA |
| | ADHD (n = 34) | 11.00 (1.77) | 17/17 | 4 Hispanic/9 White/0 Asian/1 Black/20 Other | NA |
| ABCD-site10 | TDC (n = 302) | 11.78 (1.92) | 164/138 | 114 Hispanic/84 White/6 Asian/9 Black/89 Other | NA |
| | ADHD (n = 52) | 11.39 (1.790 | 31/21 | 19 Hispanic/12 White/0 Asian/1 Black/20 Other | NA |
| ABCD-site11 | TDC (n = 143) | 11.63 (1.89) | 81/62 | 13 Hispanic/68 White/6 Asian/21 Black/35 Other | NA |
| | ADHD (n = 33) | 11.48 (1.67) | 20/13 | 4 Hispanic/12 White/2 Asian/5 Black/10 Other | NA |
| ABCD-site12 | TDC (n = 155) | 11.84 (1.89) | 74/81 | 11 Hispanic/70 White/6 Asian/38 Black/30 Other | NA |
| | ADHD (n = 41) | 11.26 (1.79) | 25/16 | 1 Hispanic/18 White/0 Asian/7 Black/15 Other | NA |
| ABCD-site13 | TDC (n = 319) | 11.97 (1.96) | 156/163 | 23 Hispanic/158 White/15 Asian/38 Black/85 Other | NA |
| | ADHD (n = 47) | 11.20 (1.81) | 28/19 | 4 Hispanic/16 White/0 Asian/4 Black/23 Other | NA |
| ABCD-site14 | TDC (n = 207) | 12.04 (1.95) | 108/99 | 20 Hispanic/137 White/2 Asian/12 Black/36 Other | NA |
| | ADHD (n = 48) | 11.41 (1.94) | 26/22 | 5 Hispanic/24 White/0 Asian/3 Black/16 Other | NA |
| ABCD-site15 | TDC (n = 160) | 11.70 (1.90) | 86/74 | 4 Hispanic/77 White/2 Asian/62 Black/15 Other | NA |
| | ADHD (n = 37) | 10.95 (1.74) | 22/15 | 1 Hispanic/11 White/0 Asian/14 Black/11 Other | NA |

| | | | | | |
|---|---|---|---|---|---|
| ABCD-site16 | TDC (n = 370) | 11.92 (1.94) | 194/176 | 31 Hispanic/248 White/7 Asian/11 Black/73 Other | NA |
| | ADHD (n = 76) | 11.32 (1.85) | 43/33 | 6 Hispanic/45 White/0 Asian/3 Black/22 Other | NA |
| ABCD-site17 | TDC (n = 175) | 11.73 (1.80) | 88/87 | 6 Hispanic/116 White/4 Asian/9 Black/40 Other | NA |
| | ADHD (n = 41) | 11.22 (1.77) | 24/17 | 2 Hispanic/24 White/0 Asian/3 Black/12 Other | NA |
| ABCD-site18 | TDC (n = 166) | 12.14 (1.96) | 90/76 | 7 Hispanic/107 White/3 Asian/8 Black/41 Other | NA |
| | ADHD (n = 24) | 11.05 (1.85) | 15/9 | 2 Hispanic/10 White/1 Asian/0 Black/11 Other | NA |
| ABCD-site19 | TDC (n = 189) | 12.36 (1.86) | 94/95 | 7 Hispanic/103 White/0 Asian/38 Black/41 Other | NA |
| | ADHD (n = 45) | 11.57 (1.84) | 28/17 | 3 Hispanic/16 White/0 Asian/8 Black/18 Other | NA |
| ABCD-site20 | TDC (n = 165) | 12.10 (1.92) | 90/75 | 4 Hispanic/86 White/1 Asian/50 Black/24 Other | NA |
| | ADHD (n = 34) | 11.37 (1.84) | 23/11 | 2 Hispanic/12 White/0 Asian/6 Black/14 Other | NA |
| ABCD-site21 | TDC (n = 189) | 11.96 (1.99) | 101/88 | 28 Hispanic/78 White/3 Asian/21 Black/59 Other | NA |
| | ADHD (n = 55) | 11.47 (1.94) | 32/23 | 5 Hispanic/16 White/1 Asian/6 Black/27 Other | NA |

#, ADHD is clinically classified into three distinct presentations on the basis of the most prominent symptoms exhibited: predominantly inattentive (ADHD-I), predominantly hyperactive/impulsive (ADHD-H/I), and combined (ADHD-C).

**Table S2. Scan parameters of the datasets**

| Dataset | Site | Vendor | Field Strength | TR/TE/TI | Flip Angle | Matrix | FOV | Slice Thickness | Slice Number |
|---|---|---|---|---|---|---|---|---|---|
| Discovery | PKU6 | GE Discovery | 3T | 6700 ms/2.90 ms/450 ms | 12 | 256×256 | 256×256 | 1 mm | 192 |
| | ADHD200-PKU | Siemens Trio | 3T | 2530 ms/3.39 ms/1100 ms | 7 | 240×256 | 240×256 | 1.33 mm | 176 |
| | | Siemens Trio | 3T | 2530 ms/3.45 ms/1100 ms | 7 | 256×256 | 256×256 | 1 mm | 176 |
| | | Siemens Trio | 3T | 2000 ms/3.67 ms/1100 ms | 12 | 256×256 | 240×240 | 1 mm | 192 |
| | | Siemens Trio | 3T | 1950 ms/2.60 ms/900 ms | 10 | 240×256 | 240×256 | 1.3 mm | 128 |
| | | Siemens Trio | 3T | 2530 ms/3.37 ms/1100 ms | 7 | 256×256 | 256×256 | 1.33 mm | 128 |
| | | Siemens Trio | 3T | 1770 ms/3.92 ms/1100 ms | 12 | 256×256 | 512×512 | 1 mm | 176 |
| | | Siemens Trio | 3T | 854 ms/2.89 ms/600 ms | 8 | 261×261 | 256×256 | 1.3 mm | 144 |
| | CCBD | Siemens Prisma | 3T | 2300 ms/2.07 ms/NA | 7 | 320×280 | 256×256 | 0.8 mm | 240 |
| | | GE Discovery | 3T | 2558 ms/3.58 ms/NA | 7 | 320×320 | 256×256 | 0.8 mm | 240 |
| | | United-Imaging | 3T | 6.5 ms/2.1 ms/NA | 7 | 320×280 | 256×256 | 0.8 mm | 240 |
| Validation | ABCD-site01 | Philips Achieva dStream | 3T | 6.31 ms/2.90 ms/1060 ms | 8 | NA | NA | 1 mm | NA |
| | ABCD-site02 | Siemens Prisma | 3T | 2500 ms/2.88 ms/1060 ms | 8 | NA | NA | 1 mm | NA |
| | ABCD-site03 | Siemens Prisma | 3T | 2500 ms/2.88 ms/1060 ms | 8 | NA | NA | 1 mm | NA |
| | ABCD-site04 | GE MR750 | 3T | 2500 ms/2.00 ms/1060 ms | 8 | NA | NA | 1 mm | NA |
| | ABCD-site05 | Siemens Prisma | 3T | 2500 ms/2.88 ms/1060 ms | 8 | NA | NA | 1 mm | NA |
| | ABCD-site06 | Siemens Prisma | 3T | 2500 ms/2.88 ms/1060 ms | 8 | NA | NA | 1 mm | NA |
| | ABCD-site07 | Siemens Prisma | 3T | 2500 ms/2.88 ms/1060 ms | 8 | NA | NA | 1 mm | NA |
| | ABCD-site08 | GE MR750 | 3T | 2500 ms/2.00 ms/1060 ms | 8 | NA | NA | 1 mm | NA |
| | ABCD-site09 | Siemens Prisma | 3T | 2500 ms/2.88 ms/1060 ms | 8 | NA | NA | 1 mm | NA |

| | | | | | | | | |
|---|---|---|---|---|---|---|---|---|
| ABCD-site10 | GE MR750 | 3T | 2500 ms/2.00 ms/1060 ms | 8 | NA | NA | 1 mm | NA |
| ABCD-site11 | Siemens Prisma | 3T | 2500 ms/2.88 ms/1060 ms | 8 | NA | NA | 1 mm | NA |
| ABCD-site12 | Siemens Prisma | 3T | 2500 ms/2.88 ms/1060 ms | 8 | NA | NA | 1 mm | NA |
| ABCD-site13 | GE MR750 | 3T | 2500 ms/2.00 ms/1060 ms | 8 | NA | NA | 1 mm | NA |
| ABCD-site14 | Siemens Prisma | 3T | 2500 ms/2.88 ms/1060 ms | 8 | NA | NA | 1 mm | NA |
| ABCD-site15 | Siemens Prisma | 3T | 2500 ms/2.88 ms/1060 ms | 8 | NA | NA | 1 mm | NA |
| ABCD-site16 | Siemens Prisma | 3T | 2500 ms/2.88 ms/1060 ms | 8 | NA | NA | 1 mm | NA |
| ABCD-site17 | Philips Achieva dStream | 3T | 6.31 ms/2.90 ms/1060 ms | 8 | NA | NA | 1 mm | NA |
| ABCD-site18 | GE MR750 | 3T | 2500 ms/2.00 ms/1060 ms | 8 | NA | NA | 1 mm | NA |
| ABCD-site19 | Philips Ingenia | 3T | 6.31 ms/2.90 ms/1060 ms | 8 | NA | NA | 1 mm | NA |
| ABCD-site20 | Siemens Prisma | 3T | 2500 ms/2.88 ms/1060 ms | 8 | NA | NA | 1 mm | NA |
| ABCD-site21 | Siemens Prisma | 3T | 2500 ms/2.88 ms/1060 ms | 8 | NA | NA | 1 mm | NA |

**Table S3. Demographic characteristics of the ADHD biotypes**

| | Infra-normal biotype | Supra-normal biotype | Statistics | P value |
|---|---|---|---|---|
| **Characteristic** | N = 238 | N = 349 | | |
| Age, years, mean (SD) | 10.5 (2.12) | 10.2 (2.15) | t = 1.69 | 0.09 |
| Sex, male/female | 185/53 | 285/64 | $\chi2$ = 1.13 | 0.29 |
| Site, PKU6/ADHD200 | 207/31 | 298/51 | $\chi2$ = 0.18 | 0.67 |
| IQ | 105.05 (16.86) | 106.90 (15.25) | t = -0.92 | 0.36 |
| **Clinical symptoms, mean (SD) #** | | | | |
| Total score | 37.90 (12.5) | 39.7 (12.5) | t = 0.44 | 0.66 |
| Hyperactive-impulsive score | 15.2 (7.95) | 16.1 (8.12) | t = 0.52 | 0.60 |
| Inattention score | 22.7 (5.85) | 23.6 (5.64) | t = 0.003 | 0.99 |
| **Comorbidity, No. subjects (%)** | | | | |
| Oppositional defiance disorder | 18 (7.56%) | 15 (4.30%) | | |
| Conduct disorder | 3 (1.26%) | 0 (%) | | |
| Learning disability | 9 (3.78%) | 2 (0.57%) | $\chi2$ = 6.47 | 0.17 |
| Tics | 7 (2.94%) | 2 (0.57%) | | |
| Others Δ | 2 (0.84%) | 3 (0.86%) | | |
| **Cognition, mean (SD) *** | | | | |
| Delay aversion (impulsivity) | 0.60 (0.26) | 0.60 (0.24) | t = -0.009 | 0.99 |
| Probability of a hit (sustained attention) | 0.72 (0.19) | 0.74 (0.18) | t = -0.69 | 0.50 |
| Stop-signal reaction time (response inhibition) | -354 (108) | -351 (118) | t = -0.23 | 0.82 |
| Between errors (spatial working memory) | -53.7 (19.1) | -49.7 (20.1) | t = -1.36 | 0.17 |
| Problems solved (spatial planning) | 5.08 (1.80) | 5.65 (2.05) | t = -1.76 | 0.08 |
| Stages completed (attention flexibility) | 7.71 (1.53) | 7.92 (1.09) | t = -0.98 | 0.33 |

### Data available for 508 participants (infra-normal biotype = 211, supra-normal biotype = 297)
*Data available for 164 participants (infra-normal biotype = 49, supra-normal biotype = 115)
Δ Others: social anxiety disorder, depression, and social phobia.

Data are presented as the mean (SD) or n (%). The ADHD200 cohort included only the PKU site. TDC, typically developing children; ADHD, attention-deficit/hyperactivity disorder.

#### Table S4. Pre- and posttreatment scores and drug types in each ADHD biotype

| Biotype (sample size) | Drug | Total score | | Hyperactivity-impulsivity score | | Inattention score | |
|---|---|---|---|---|---|---|---|
| | | Pretreatment | Posttreatment | Pretreatment | Posttreatment | Pretreatment | Posttreatment |
| Infra-normal biotype (N = 34) | MPH | 29.29 (6.74) | 18.06 (10.06) | 11.47 (5.91) | 6.50 (5.29) | 17.82 (3.67) | 11.56 (5.58) |
| Supranormal biotype (N = 50) | MPH | 29.08 (9.11) | 17.08 (8.19) | 11.10 (6.80) | 5.52 (4.41) | 17.98 (4.15) | 11.56 (5.11) |
| Infra-normal biotype (N = 15) | ATX | 28.87 (6.69) | 23.73 (8.15) | 10.93 (5.64) | 9.27 (5.81) | 17.93 (3.63) | 14.47 (3.70) |
| Supranormal biotype (N = 29) | ATX | 30.52 (9.55) | 19.44 (10.02) | 11.83 (7.28) | 7.93 (5.52) | 18.69 (3.59) | 11.52 (5.23) |

Scores are mean (SD). MPH: methylphenidate, ATX: atomoxetine. The children with ADHD were selected from the PKU6 cohort.

**Table S5. Post hoc analysis of two-way repeated-measures ANOVA for total symptom score**

| Factor | F value (Df) | P value |
|---|---|---|
| ***Infra-normal biotype*** | | |
| Time | 59.86 (1, 47) | **<0.001** |
| Medication type | 1.43 (1, 47) | 0.24 |
| Time*Medication type | 5.40 (1, 47) | **0.02** |
| **Post hoc: MPH** | | |
| Time | 51.20 (1, 33) | **<0.001** |
| **Post hoc: ATX** | | |
| Time | 9.08 (1, 14) | **<0.001** |
| ***Supranormal biotype*** | | |
| Time | 127.83 (1, 77) | **<0.001** |
| Medication type | 1.08 (1, 77) | 0.30 |
| Time*Medication type | 0.19 (1, 77) | 0.67 |
| **Post hoc: MPH** | | |
| Time | 74.02 (1, 49) | **<0.001** |
| **Post hoc: ATX** | | |
| Time | 58.46 (1, 28) | **<0.001** |

MPH: methylphenidate; ATX: atomoxetine; ANOVA, analysis of variance.

**Table S6. Post hoc analysis of two-way repeated-measures ANOVA for hyperactivity-impulsivity scores**

| Factor | F value (Df) | P value |
|---|---|---|
| ***Infra-normal biotype*** | | |
| Time | 30.18 (1, 47) | **<0.001** |
| Medication type | 0.51 (1, 47) | 0.48 |
| Time*Medication type | 4.46 (1, 47) | **0.04** |
| **Post hoc: MPH** | | |
| Time | 29.52 (1, 33) | **<0.001** |
| **Post hoc: ATX** | | |
| Time | 2.27 (1, 14) | 0.15 |
| ***Supranormal biotype*** | | |
| Time | 71.29 (1, 77) | **<0.001** |
| Medication type | 1.53 (1, 77) | 0.22 |
| Time*Medication type | 1.91 (1, 77) | 0.17 |
| **Post hoc: MPH** | | |
| Time | 55.43 (1, 49) | **<0.001** |
| **Post hoc: ATX** | | |
| Time | 17.01 (1, 28) | **<0.001** |

MPH: methylphenidate, ATX: atomoxetine, ANOVA, analysis of variance.

**Table S7. Post hoc analysis of two-way repeated-measures ANOVA for inattention score**

| Factor | F value (Df) | P value |
|---|---|---|
| ***Infra-normal biotype*** | | |
| Time | 40.23 (1, 47) | **<0.001** |
| Medication type | 2.19 (1, 47) | 0.15 |
| Time*Medication type | 2.30 (1, 47) | 0.14 |
| **Post hoc: MPH** | | |
| Time | 34.82 (1, 33) | **0.001** |
| **Post hoc: ATX** | | |
| Time | 6.28 (1, 14) | **0.03** |
| ***Supranormal biotype*** | | |
| Time | 120.25 (1, 77) | **<0.001** |
| Medication type | 0.15 (1, 77) | 0.70 |
| Time*Medication type | 0.35 (1, 77) | 0.55 |
| **Post hoc: MPH** | | |
| Time | 59.32 (1, 49) | **<0.001** |
| **Post hoc: ATX** | | |
| Medication type | 73.78 (1, 28) | **<0.001** |

MPH: methylphenidate, ATX: atomoxetine, ANOVA, analysis of variance.

#### Table S8. ADHD GWAS genes

| |
|---|
| ABHD5 |
| AC005493.1 |
| AC018797.2 |
| AC084125.2 |
| ADAMTS17 |
| AL024506.1 |
| ANO10 |
| AP006621.1 |
| ARHGAP39 |
| ARTN |
| ASTN2 |
| ATP11A |
| ATP2C2 |
| ATP6V0B |
| ATP6V1F |
| B4GALT2 |
| BCHE |
| BMPR1B |
| BSN |
| C10orf79 |
| C12orf68 |
| C1orf173 |
| C21orf34 |
| C8orf82 |
| C9orf98 |
| CADM2 |
| CALD1 |
| CALN1 |
| CAMKV |
| CARF |
| CCDC136 |
| CCDC24 |
| CCDC46 |
| CCDC71 |
| CDC20 |
| CDH13 |
| CDH23 |
| CDH8 |
| CHRNA7 |

CNBD1
CNTN2
CNTNAP2
CNTNAP5
COL19A1
COPA
CPSF2
CREB5
CRYGC
CSMD1
CSMD2
CTC-498M16.4
CTD-2330K9.3
CTNNA2
CUBN
DCC
DMRT2
DMRTB1
DNM1
DPH2
DUSP1
DUSP6
DYNC2H1
ELOVL1
ELOVL6
EMP2
EPHA7
FAM190A
FAM5B
FHIT
FLNC
FOXP1
FOXP2
GABRB2
GFOD1
GIGYF2
GNAT1
GPC6
GRIK1
GRIK4

GRM1

GRM3

GRM5

GRM7

GRM8

HK1

HKDC1

HYAL3

HYI

IL16

IPO13

IQCF1

IQCF2

IQCF3

IQCF5

ITGA1

ITGA11

ITGAE

ITPR2

KCNC1

KCNIP4

KCNJ13

KCP

KCTD16

KDM4A

KDM4A-AS1

KIZ

KLHL1

LECT1

LINC00461

LINC00951

LINC01288

LINC01572

LINC02060

LINC02497

LRFN2

LRP1B

LSM6

MAML3

MAN2A2

MANBA
MAP1B
MAPK1
MBOAT1
MCTP1
MED8
MEF2C
MEIS2
METTL15
MIR3666
MIR9-2
MMP24
MOBP
MON1A
MPL
MSH3
MST1R
MTA3
MYC
MYT1L
NAP5
NAPRT1
NBEAL1
NCSTN
NDUFA5
NDUFB1
NEDD4L
NFASC
NKX2-4
NOS1
NRXN1
NT5DC3
NUCB1
NXPH1
OSBPL11
PARK2
PCDH15
PCDH7
PCDP1
PDCD6IP

PEA15
PICK1
PIDD1
PLOD3
PNPLA2
POC1B
POLR1A
POLR2B
POLR3C
PPM1F
PPM1H
PPP1R16A
PREX2
PRKCQ
PRKG1
PSD3
PTPRF
RBFOX1
RBMS3
REEP5
RNF219-AS1
ROBO2
RORA
RP11-6L6.2
RPLP2
SCN10A
SCN11A
SEMA3F
SEMA6D
SEP5
SH3BGRL2
SLC10A7
SLC2A3
SLC38A1
SLC6A9
SLCO3A1
SNRK
SOBP
SORCS3
SPAG16

| |
|---|
| SPOCK3 |
| ST3GAL3 |
| STGAL3 |
| STK32C |
| SUPT3H |
| SV2C |
| SYT10 |
| SZT2 |
| TBL1XR1 |
| TCERG1L |
| TCF12 |
| TIE1 |
| TLL2 |
| TMEM161B-AS1 |
| TMEM200C |
| TRAIP |
| TRIM32 |
| TSC22D2 |
| TTC27 |
| TUBA3C |
| TUBGCP2 |
| UGT1A9 |
| UNC5B |
| VGLL3 |
| WASL |
| WDR12 |
| WWOX |
| XRN2 |
| ZMIZ1 |
| ZNF385D |
| ZNF423 |
| ZNF544 |

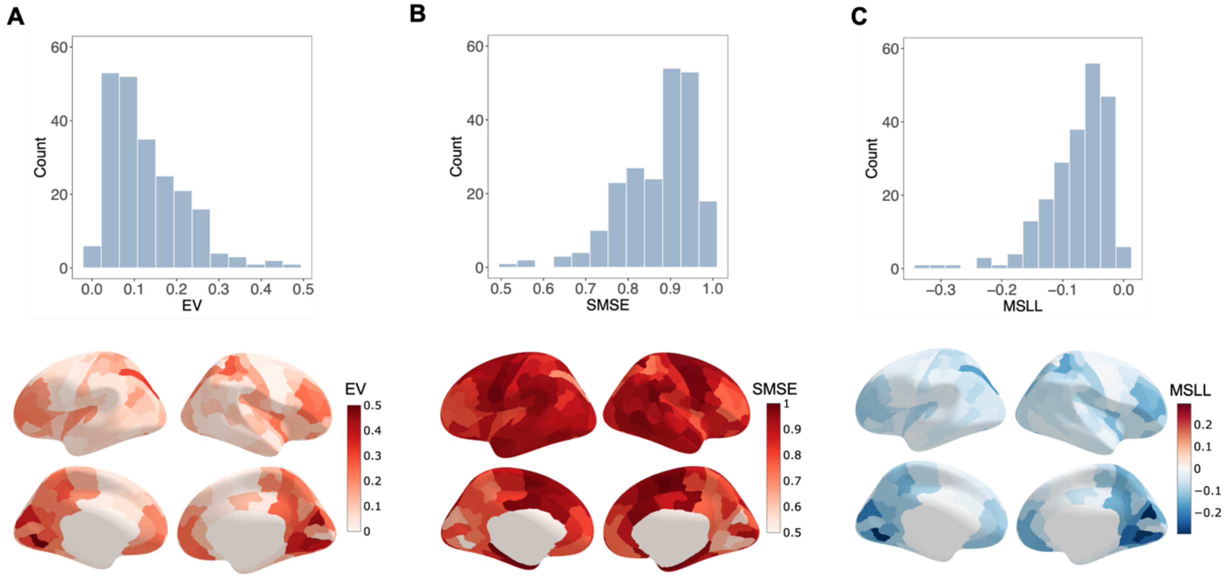


**Figure S1. Normative model performance. (A)** Distributions of explained variance (EV). **(B)** Standardized mean squared error (SMSE). **(C)** Mean standardized log-loss (MSLL). The upper histogram shows the distribution of each metric across 219 cortical regions, and the lower brain map illustrates the spatial distribution of each metric.

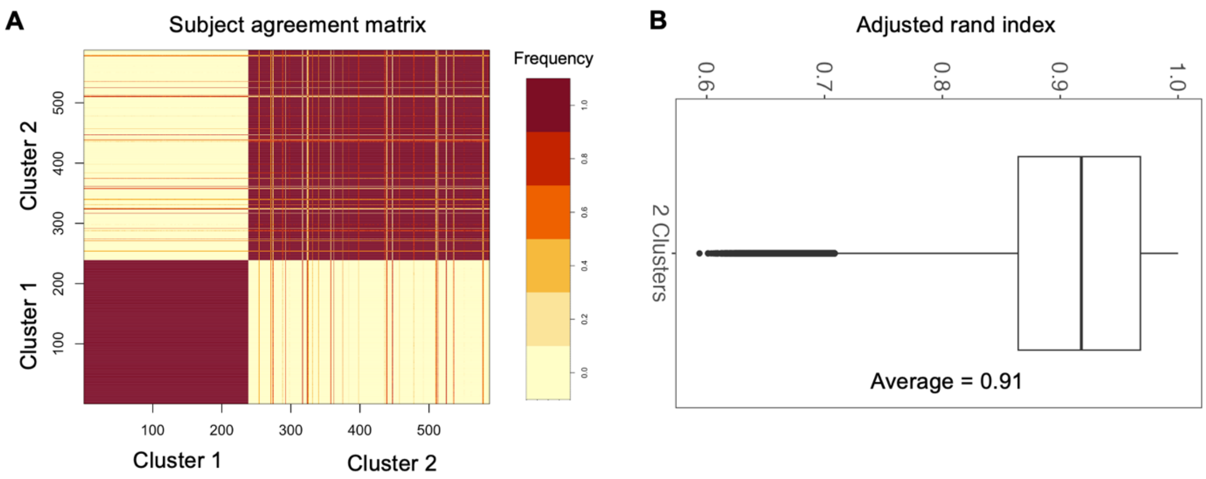


**Figure S2. Stability of clustering assessed by bootstrapping. (A)** The participant agreement matrix illustrates how often each participant clustered with another participant across 1,000 resamplings. **(B)** Adjusted rand index (ranging from 0 to 1), which measures the overlap between clusters across 1,000 resamplings. A value of 0 indicates completely random clusters across 1,000 resamplings, and a value of 1 indicates identical clusters across resamplings. The adjusted rand index of 0.91 indicated a very similar clustering solution across resamplings.

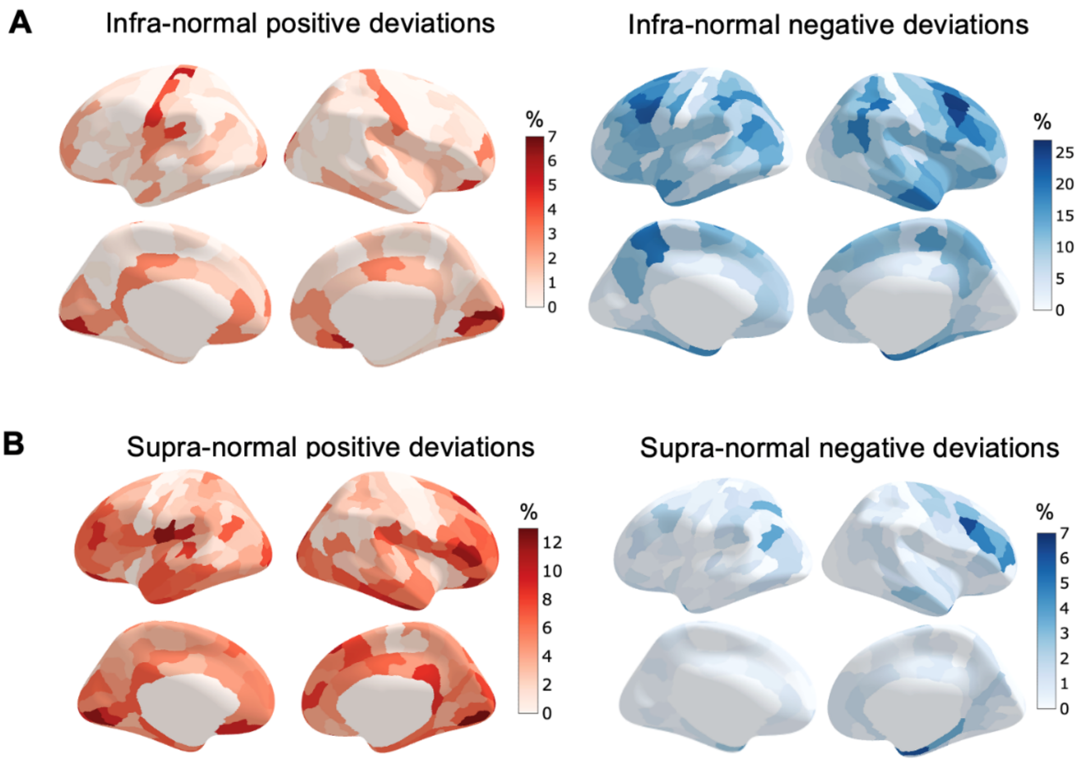


**Figure S3. Extreme deviations of cortical thickness in each ADHD biotype. (A, B)** Spatial maps illustrating the percentage of individuals within the infra-normal **(A)** and supra-normal **(B)** biotypes with extreme deviations (|Z| > 2) in each region. Positive and negative deviations are displayed on the left and right, respectively.

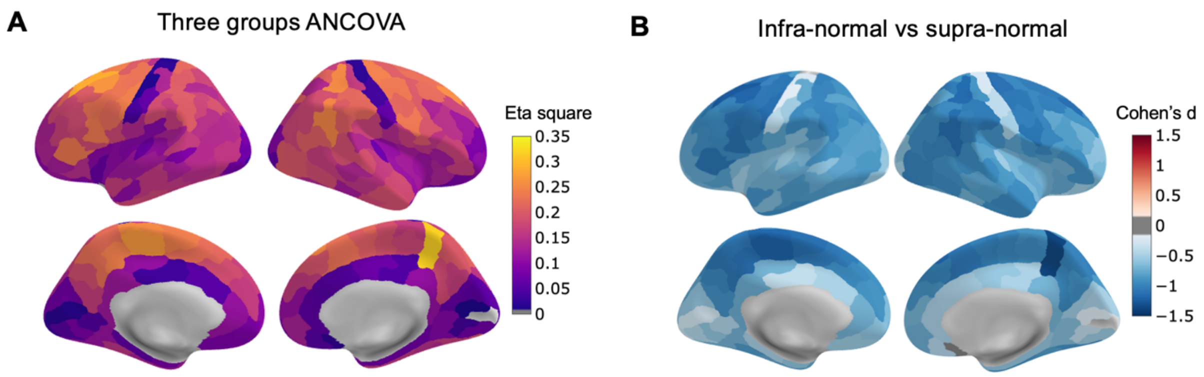


**Figure S4. Case–control analysis of cortical thickness in the infra-normal, supra-normal, and control groups**. **(A)** One-way analysis of covariance (ANCOVA) of cortical thickness among the two ADHD biotypes and the testing controls. **(B)** Post hoc results showing differences between the two biotypes. All analyses included age, sex, and drug as covariates, and multiple comparisons were corrected using a false discovery rate (FDR-corrected $q < 0.05$).

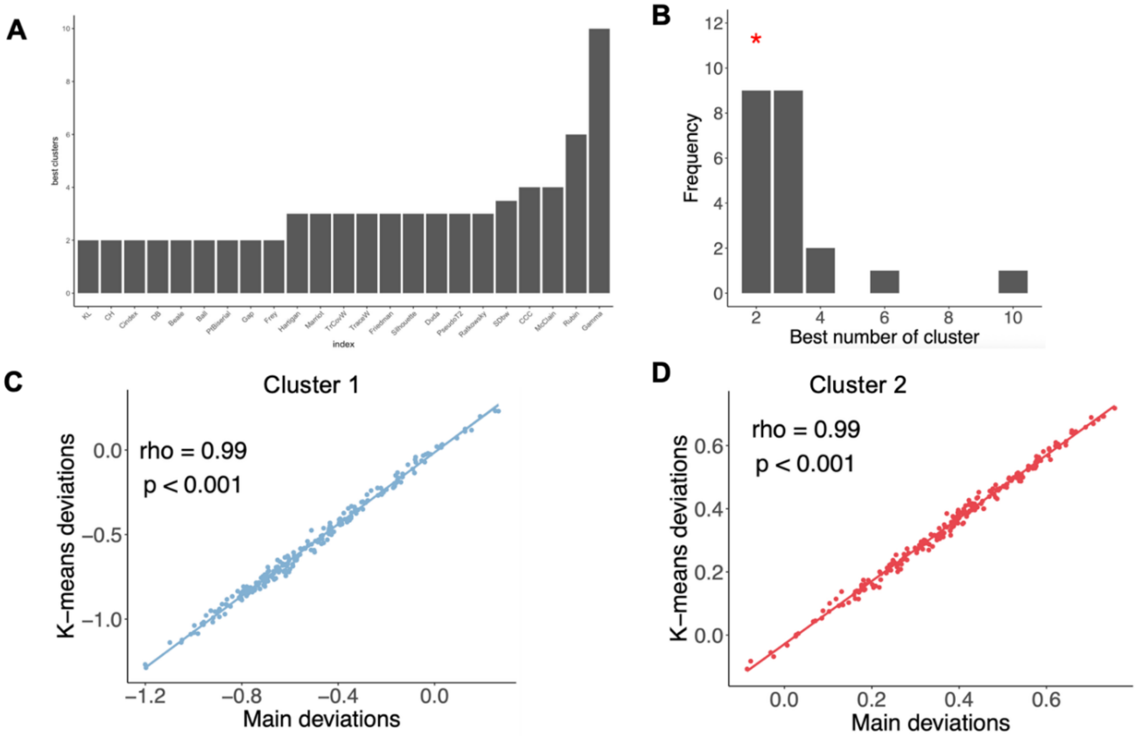


**Figure S5. Validation of biotype stability using K-means clustering. (A)** Evaluation of cluster solutions ranging from 2 to 10. **(B)** The optimal number of clusters (n = 2) was determined by a winner-take-all approach across 30 indices (NbClust). **(C, D)** Spatial correlation of average cortical deviations between the primary spectral clustering and K-means clustering solutions for the infra-normal (**C**) and supra-normal (**D**) biotypes.

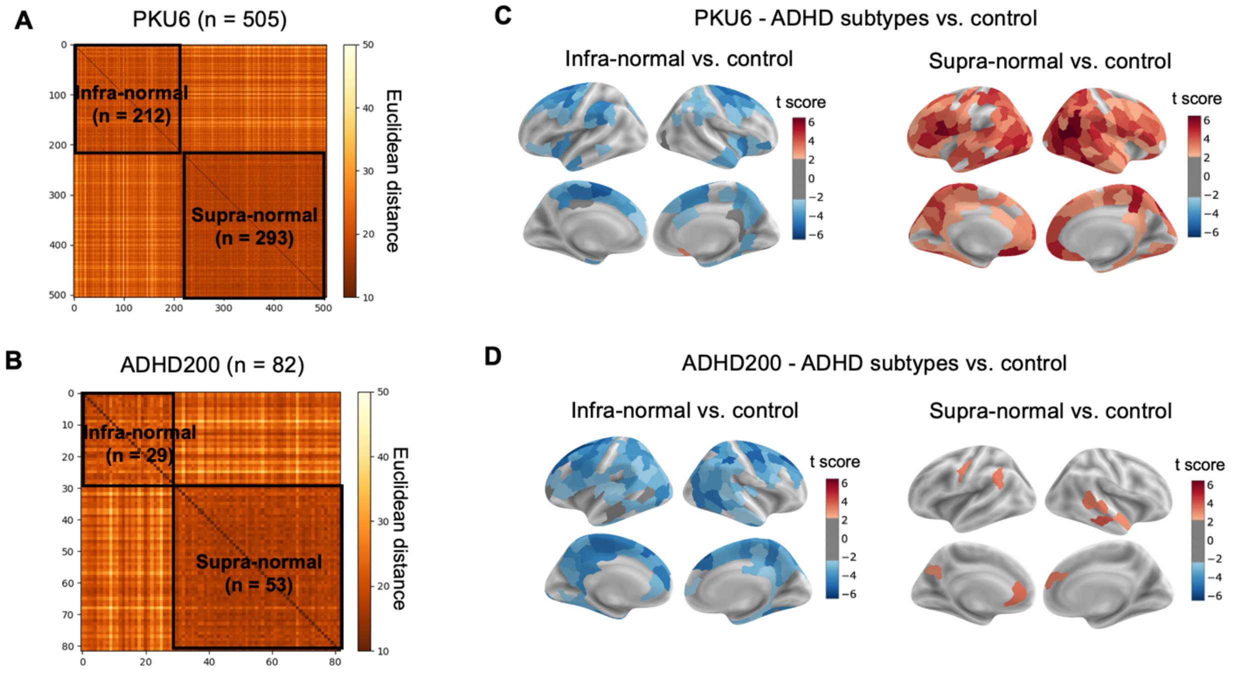


**Figure S6. Consistent identification of two ADHD biotypes across independent sites. (A, B)** Intersubject similarity matrices of cortical deviations for the PKU6 (**A**) and ADHD200 (**B**) sites. Independent clustering within each site consistently identified two distinct biotypes. (**C, D**) Case–control differences in cortical thickness (*T*-maps) for each ADHD biotype at the PKU6 **(C)** and ADHD200 **(D)** sites ($q < 0.05$, FDR corrected).

A

| Symptoms | Z | P |
|---|---|---|
| Inattention | 66.07 | <0.001 |
| Hyperactivity-impulsivity | 95.55 | <0.001 |
| Total | 3741.8 | <0.001 |

B

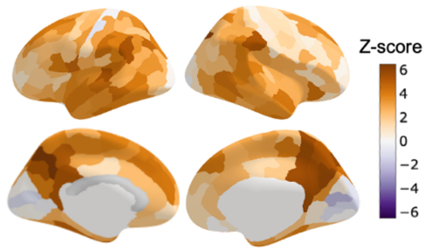


C

| Symptoms | Z | P |
|---|---|---|
| Inattention | 69.07 | <0.001 |
| Hyperactivity-impulsivity | 84.10 | <0.001 |
| Total | 1778.5 | <0.001 |

D

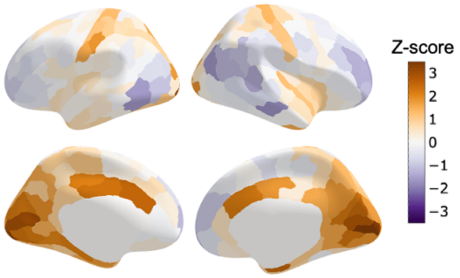


E

| Cognition | Z | P |
|---|---|---|
| Delay aversion | 11.84 | <0.001 |
| Sustained attention | -9.98 | <0.001 |
| Response inhibition | -5.32 | <0.001 |
| Spatial working memory | -13.24 | <0.001 |
| Spatial planning | -24.58 | <0.001 |
| Attention flexibility | -6.79 | <0.001 |

F

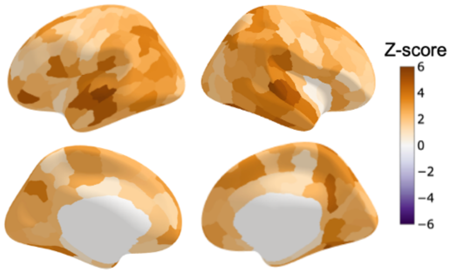


**Figure S7. Stability of contribution weights assessed via bootstrapping resampling. (A–D)** Symptom weights and their corresponding brain weights for the infra-normal **(A, B)** and supra-normal **(C, D)** biotypes. **(E, F)** Cognitive weights (**E**) and their corresponding brain weights (**F**) for the supra-normal biotype.

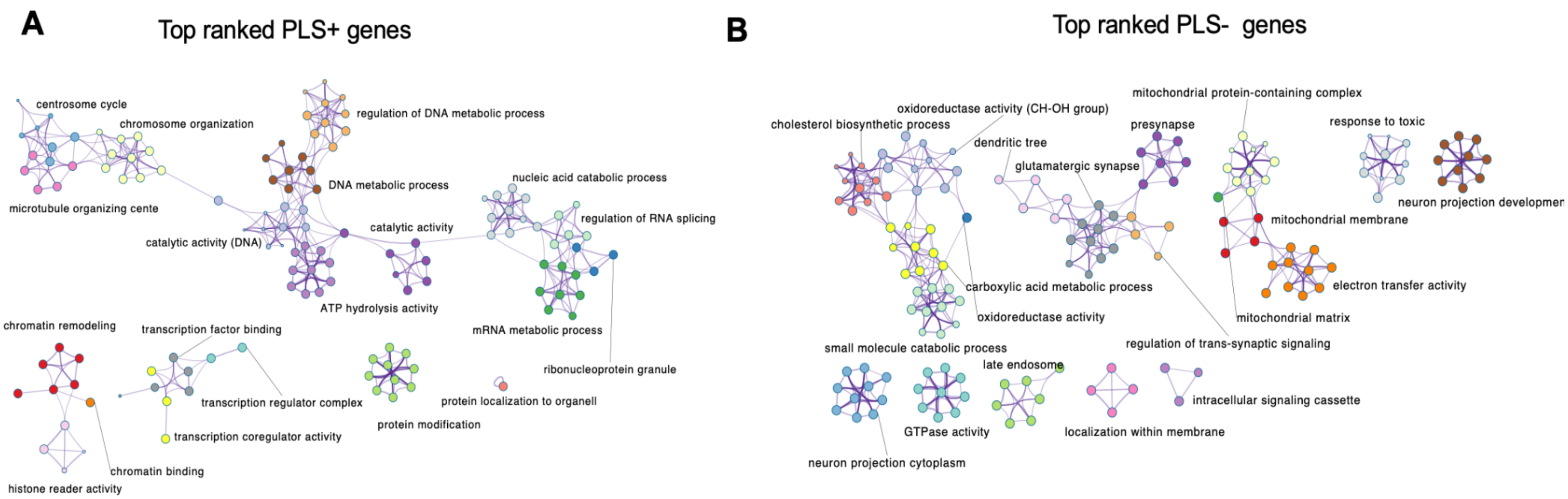


**Figure S8. Functional enrichment networks for the infra-normal biotype. (A, B)** Enrichment networks for the top-ranked PLS+ (**A**) and PLS− (**B**) genes. The network layout illustrates within- and between-cluster similarities of the enriched terms. The nodes represent individual terms, with color indicating cluster identity and size reflecting the number of input genes. The edges denote the similarity values between terms.

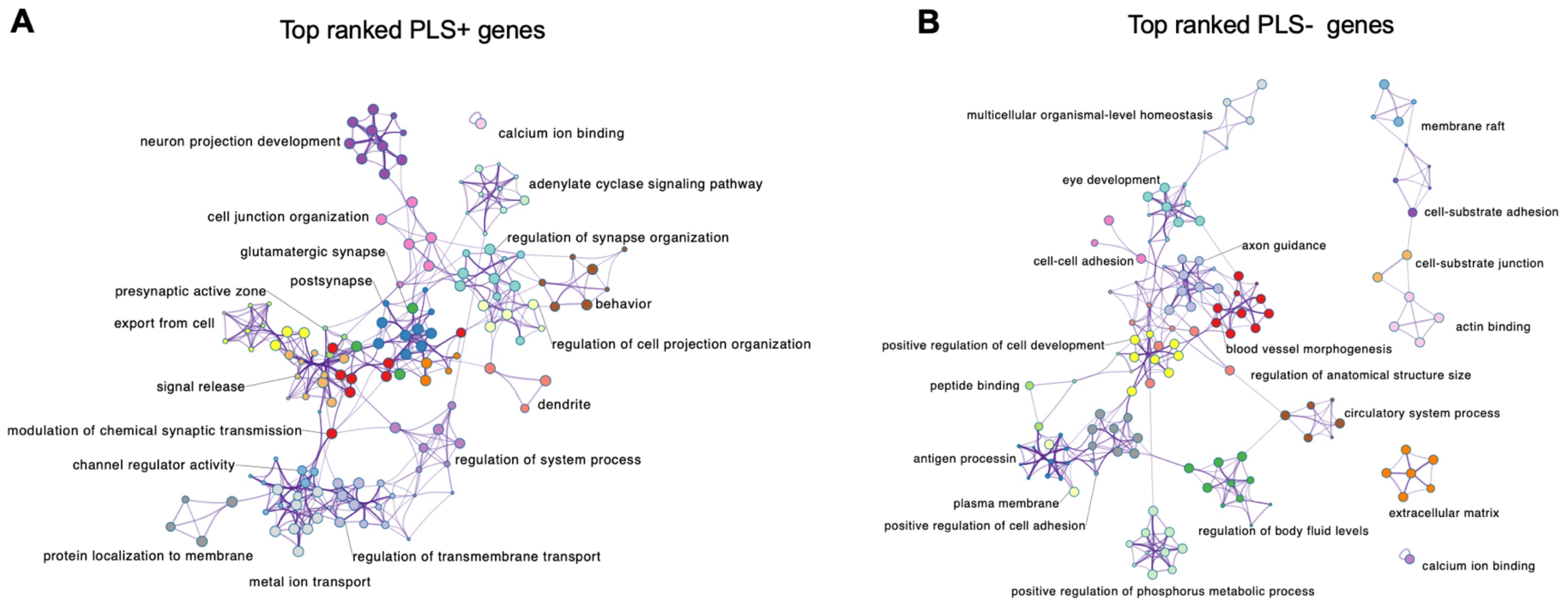


**Figure S9. Functional enrichment networks for the supra-normal biotype. (A, B)** Enrichment networks for the top-ranked PLS+ (**A**) and PLS− (**B**) genes. The network layout illustrates within- and between-cluster similarities of the enriched terms. The nodes represent individual terms, with color indicating cluster identity and size reflecting the number of input genes. The edges denote the similarity values between terms.

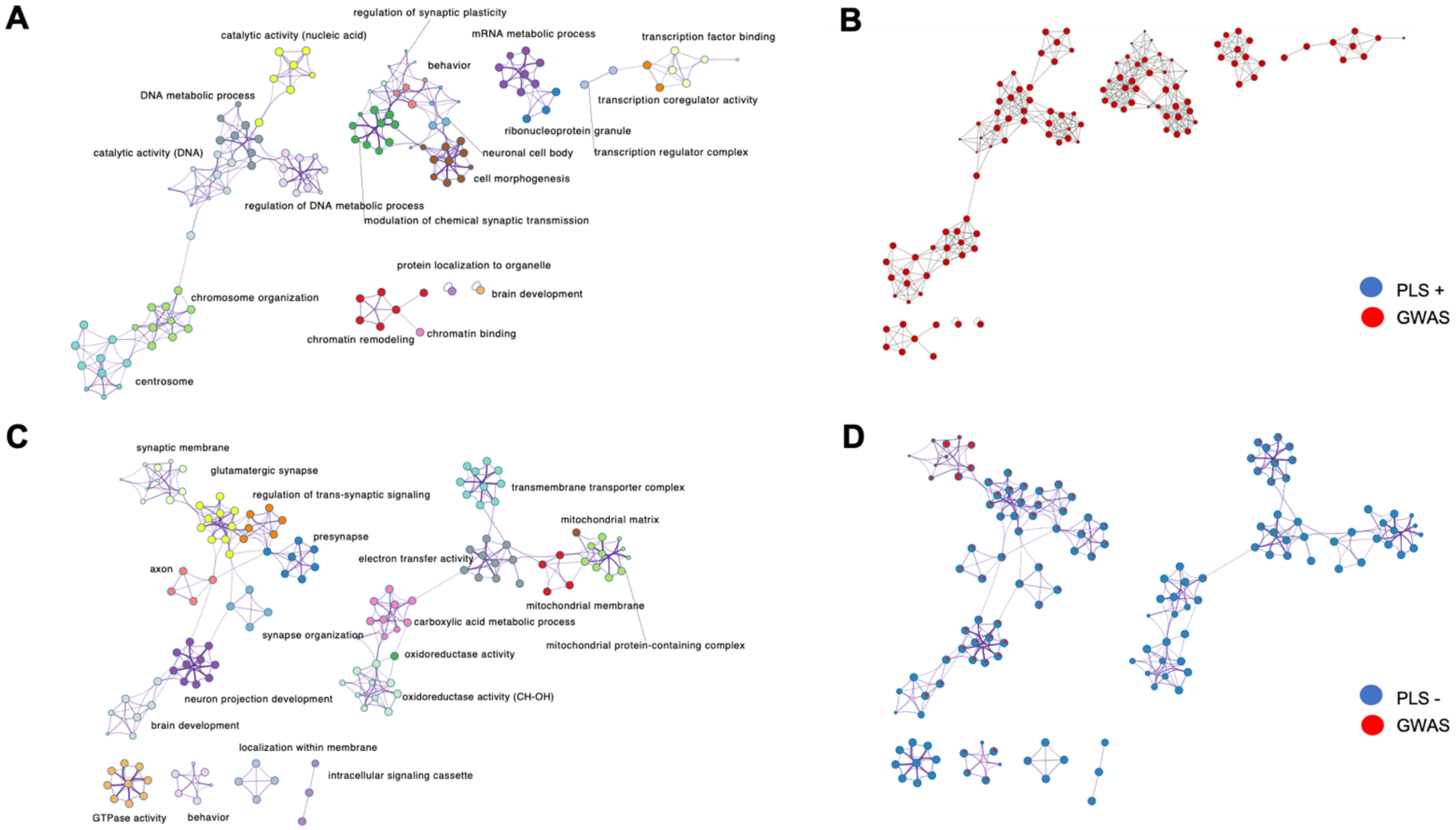


**Figure S10. Shared functional enrichment between infra-normal biotype PLS genes and ADHD GWAS polygenic risk genes.** (**A**) Enrichment network of terms associated with PLS+ and ADHD GWAS genes. The node colors represent distinct functional clusters. (**B**) The same network with nodes displayed as pie charts, illustrating the relative contribution of each gene list to a given term (blue: PLS+ genes; red: ADHD GWAS genes). (**C, D**) Corresponding enrichment networks for PLS− and ADHD GWAS genes, formatted as in A and B.

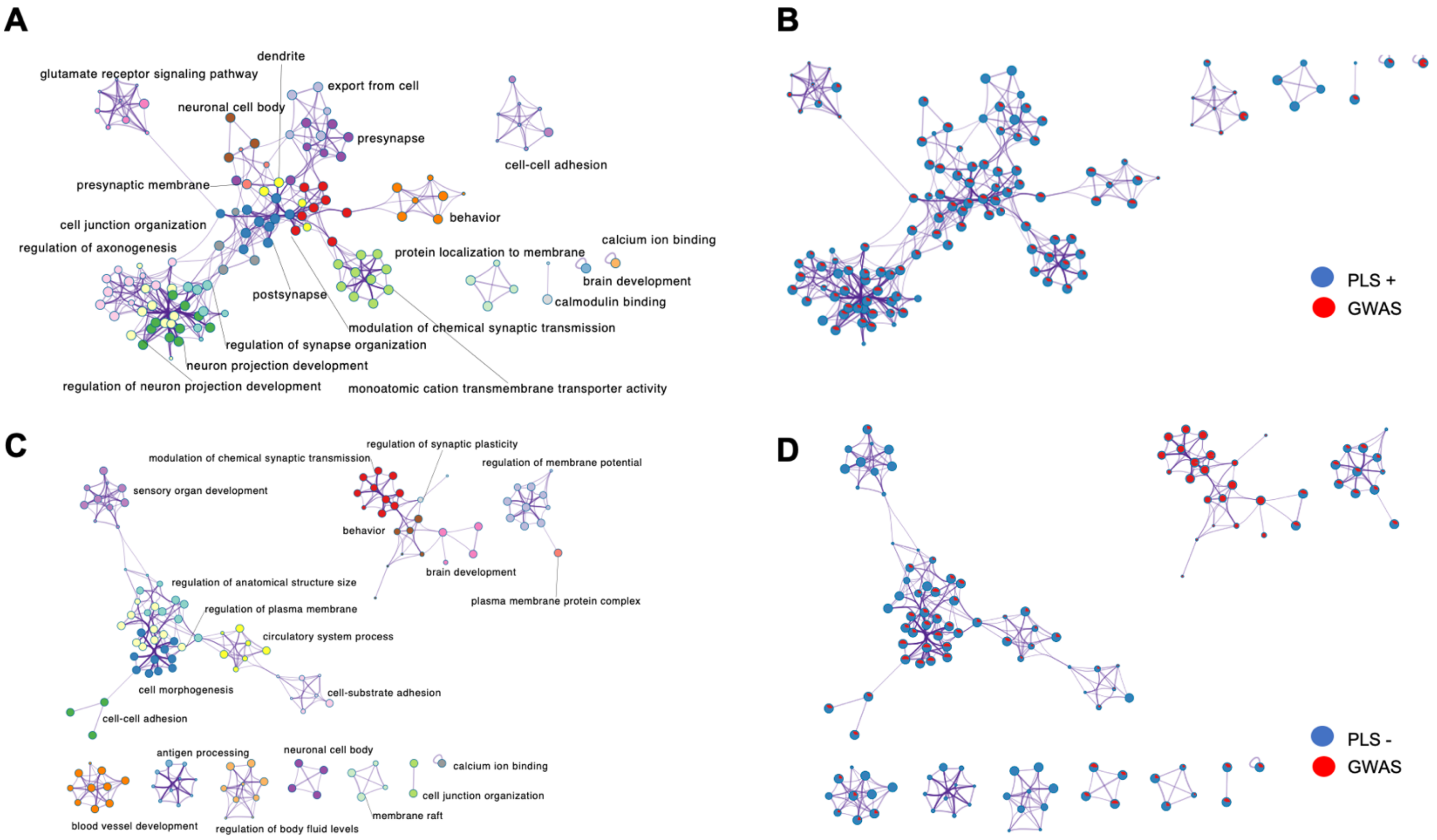


**Figure S11. Shared functional enrichment between supra-normal biotype PLS genes and ADHD GWAS polygenic risk genes.** (**A**) Enrichment network of terms associated with PLS+ and ADHD GWAS genes. The node colors represent distinct functional clusters. (**B**) The same network with nodes displayed as pie charts, illustrating the relative contribution of each gene list to a given term (blue: PLS+ genes; red: ADHD GWAS genes). (**C, D**) Corresponding enrichment networks for PLS− and ADHD GWAS genes, formatted as in A and B.

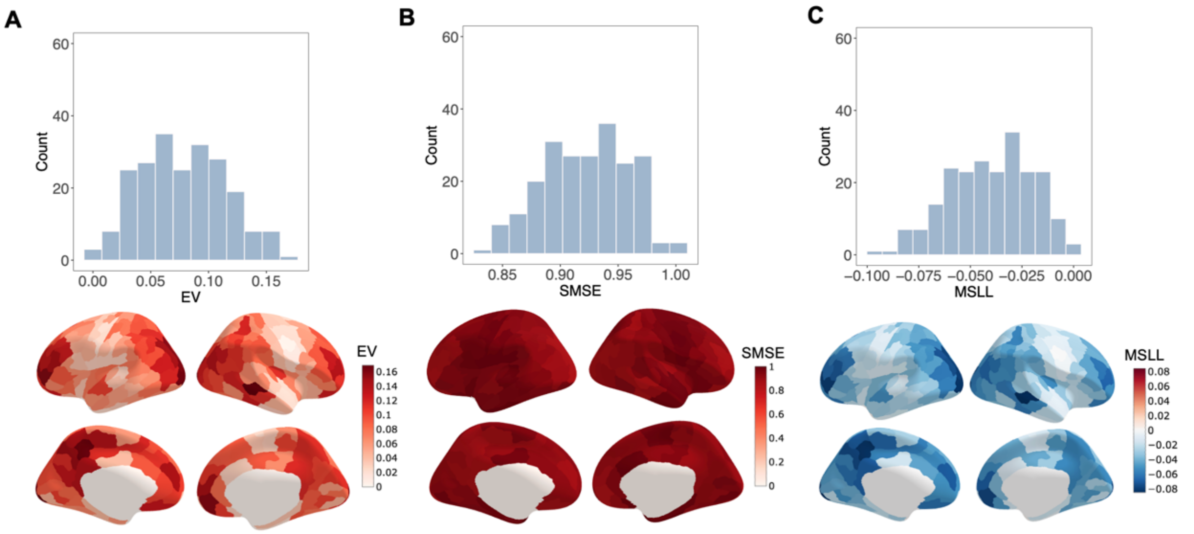


**Figure S12. Normative model performance in the ABCD cohort. (A–C)** Model performance evaluated via explained variance (EV; **A**), standardized mean squared error (SMSE; **B**) and mean standardized log-loss (MSLL; **C**). For each metric, the top histogram displays its distribution across 219 cortical regions, and the bottom brain maps illustrate its spatial topography.

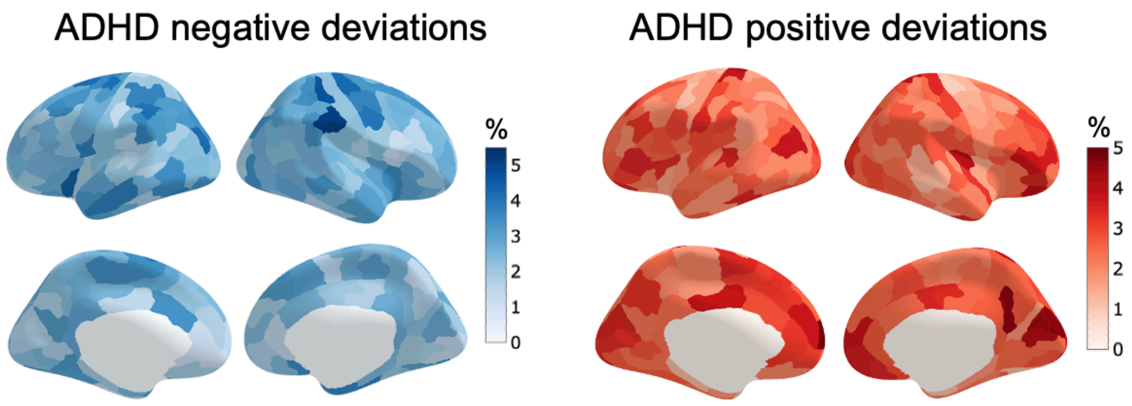


**Figure S13. Extreme deviations in cortical thickness in the ADHD group within the ABCD cohort.** Spatial distribution maps illustrating the percentage of individuals with ADHD with extreme positive and negative deviations (|Z| < 2) in each region.

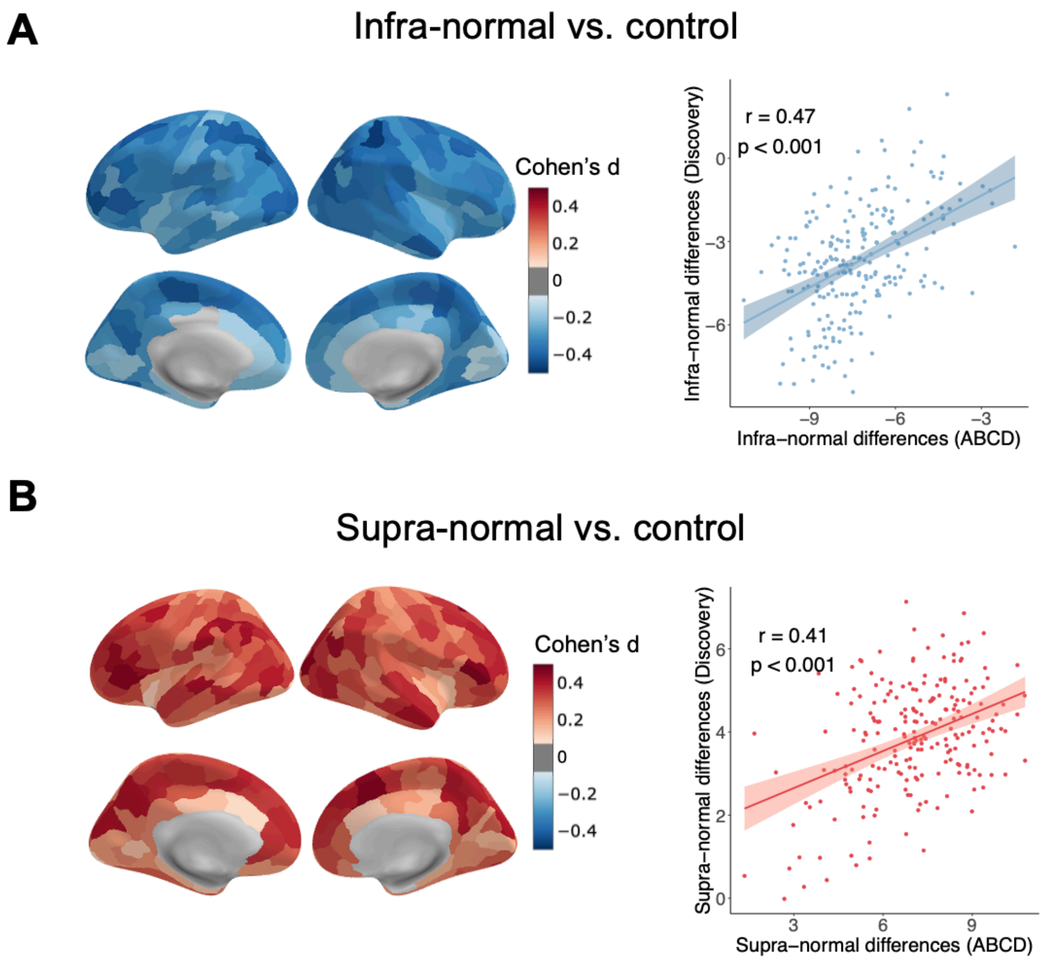


**Figure S14. Differences in cortical thickness between each ADHD biotype and controls in the ABCD cohort. (A, B)** Group differences in cortical deviations between controls and the infra-normal (**A**) or supra-normal (**B**) biotypes (left panels). Scatter plots (right panels) illustrate the spatial correlations (spin tests) of these deviation patterns between the discovery and validation datasets.
